# Versatile high-speed volumetric imaging from microscopic to macroscopic scale by self-adaptive oblique plane microscopy

**DOI:** 10.64898/2025.12.16.694458

**Authors:** Dominique Meyer, Grant Kroeschell, Xiankun Lu, Linh Hoang, Haochen Wang, Shuying Li, Yu Kang T. Xu, Lei Tian, Jeff S. Mumm, Dwight E. Bergles, Ji Yi

**Author notes:** These authors contributed equally (Dominique Meyer, Grant Kroeschell, Xiankun Lu, Linh Hoang, Ji Yi).

## Abstract

There is an increasing need for large-scale high-speed volumetric recording in complex multi-cellular model systems to define dynamic processes. Oblique plane microscopy (OPM) provides a solution that features oblique illumination, rapid optical scanning, and remote focusing to achieve real-time 4D microscopy. OPM implements light sheet imaging via a single primary objective lens, making the entire space below the objective accessible for large specimens, such as living mouse brain. Yet it is challenging to adopt OPM beyond a microscopic scale (*i.e.* size < 1mm), limiting its broad applications. Here we present a self-adaptive OPM that leverages Abbe’s sine condition to unlock its flexibility across a range of field-of-views (FOVs) (up to 8 mm^2^) and resolutions (down to 2.2 µm^3^). This versatility enables brain-wide single neuron volumetric calcium imaging in behaving larval zebrafish (1×0.4 mm^2^ FOV at 5 Hz) and capillary blood cell tracking in living mouse brain (>3×3 mm^2^ FOV) with a sweeping 0.32 mm wide volume section at 100 Hz. In optically cleared mouse brain, the flexibility allows a screen-and-zoom capability by sequentially imaging the whole brain at low-and-high magnifications to locate and resolve subcellular structures such as dendritic tress and spines. By offering a switchable imaging resolution, volume, and speed, the self-adaptive OPM achieves a versatile platform for studying a wide range of multi-cellular model system, whether *in vivo* or fixed and optically cleared.

## Introduction

Light sheet fluorescence microscopy (LSFM) is a transformative imaging method in the fields of developmental biology^1^, neuroscience^2^, and pathology^3^. The high illumination efficiency, low photobleaching, and low phototoxicity of this epifluorescence modality coupled with a sensitive scientific camera enables high-throughput volumetric imaging. Recently, oblique plane microscopy (OPM), a type of LSFM, has attracted considerable attention by employing one primary objective for both excitation and emission pathways, alleviating the complex dual-objective alignment and the sample space constraints in conventional light sheet systems^4,5,6^. Further, by combining rapid optical scanning with a high-speed camera, the volumetric imaging speed can achieve up to 300 volumes per second^6^. These features have established high-speed OPM as an optimal tool to image across a range of dynamic biological applications such as vascular flow and calcium imaging of the beating embryonic zebrafish heart^7^, embryonic egg chamber development in *Drosophila*^8^, and whole-brain activity in zebrafish larvae^8^.

However, high-speed OPM is limited by a field-of-view (FOV) of only several hundred microns, making it unsuitable for mesoscopic or macroscopic imaging applications that require millimeter-scale volumetric ranges. There is high demand for this capability, as cell signaling events within tissues and organs often propagate over these distances. The prevailing design of OPM optics uses the principle of perfect remote imaging via unity magnification between object space and intermediate imaging space (IIS) to eliminate spherical aberration^9^. The oblique light sheet is de-scanned and reimaged at the IIS, and the tertiary remote imaging system is rotated to refocus the oblique light sheet on the planar detector. The rotation of the remote imaging system crops the primary numerical aperture (NA), and the light loss can be absolute beyond a threshold angle of rotation^10^. While the index mismatched method can minimize light loss by using the full NA of the primary objective^11^, it requires a large oblique light sheet angle provided by a high NA objective with high magnification, which comes at the cost of FOV.

There are different approaches to address this limitation by manipulating the oblique angle either proximally on the sample side, or distally at the IIS. At the proximal end, the illumination angle can be increased through an additional diffraction grating^12^, microprism^13^, or external light sheet illumination independent of the primary objective^14,15^. At the distal end, several approaches have been proposed to re-direct the light into the remote imaging system using an index-mismatched interface^16^, diffraction grating^17^, fiber optic faceplate^18^, or bespoke objective lens^19^. Each of these solutions increases system complexity and needs to be tailored for specific imaging parameters (e.g. resolution, FOV), limiting their practical accessibility and widespread adoption.

In this study, we present a simple design principle that achieves self-adaptation in OPM from microscopic to macroscopic imaging, prioritizing FOV over absolute optical perfection. Our approach leverages Abbe’s sine condition, which expands OPM imaging to accommodate different primary objectives across varying FOVs, resolutions, and magnifications without requiring additional optical components. We demonstrate this capability using a 4× objective for a wide-field view (FOV 8×8 mm^2^, resolution 6×7×60 µm^3^) to visualize global patterns, a 10× objective to image individual cells (FOV 3.2×3.2 mm^2^, resolution 3×3×10 µm^3^ x-y-z), and a 20× objective to achieve sub-cellular resolution (FOV 1.5×1.0 mm^2^, resolution 1.2×2×2.2 µm^3^ x-y-z), all within the same experimental platform. We allowed modest aberrations inherent to this multi-objective approach, achieving a comprehensive imaging platform that bridges the traditional gap between macroscopic, mesoscopic, and high-resolution subcellular imaging. We show that this method achieves high performance across diverse imaging contexts: resolving individual dendritic projections in cleared whole mouse brain samples, exploring sensorimotor neural activity patterns in larval zebrafish, and mapping vascular networks spanning large vessels to single capillaries in the mouse brain *in vivo.* OPM brings the familiar versatility of conventional microscopy to advanced light-sheet applications while significantly expanding the experimental scope for the broader scientific community.

## Results

### 1. Self-adaptive optics and flexibility from microscopic to macroscopic imaging by OPM

**Figure 1a** shows the general system schematic (detailed rendering in **Supplemental Fig. S1)**, following a general configuration of OPM but incorporating a switchable primary objective lens. The light sheet illumination optics are mounted on a platform to allow slight adjustments of the illumination angle. The preferred resolution and FOV can be chosen at will by selecting primary objective lenses with desired magnifications and resolutions.

**Fig. 1:**
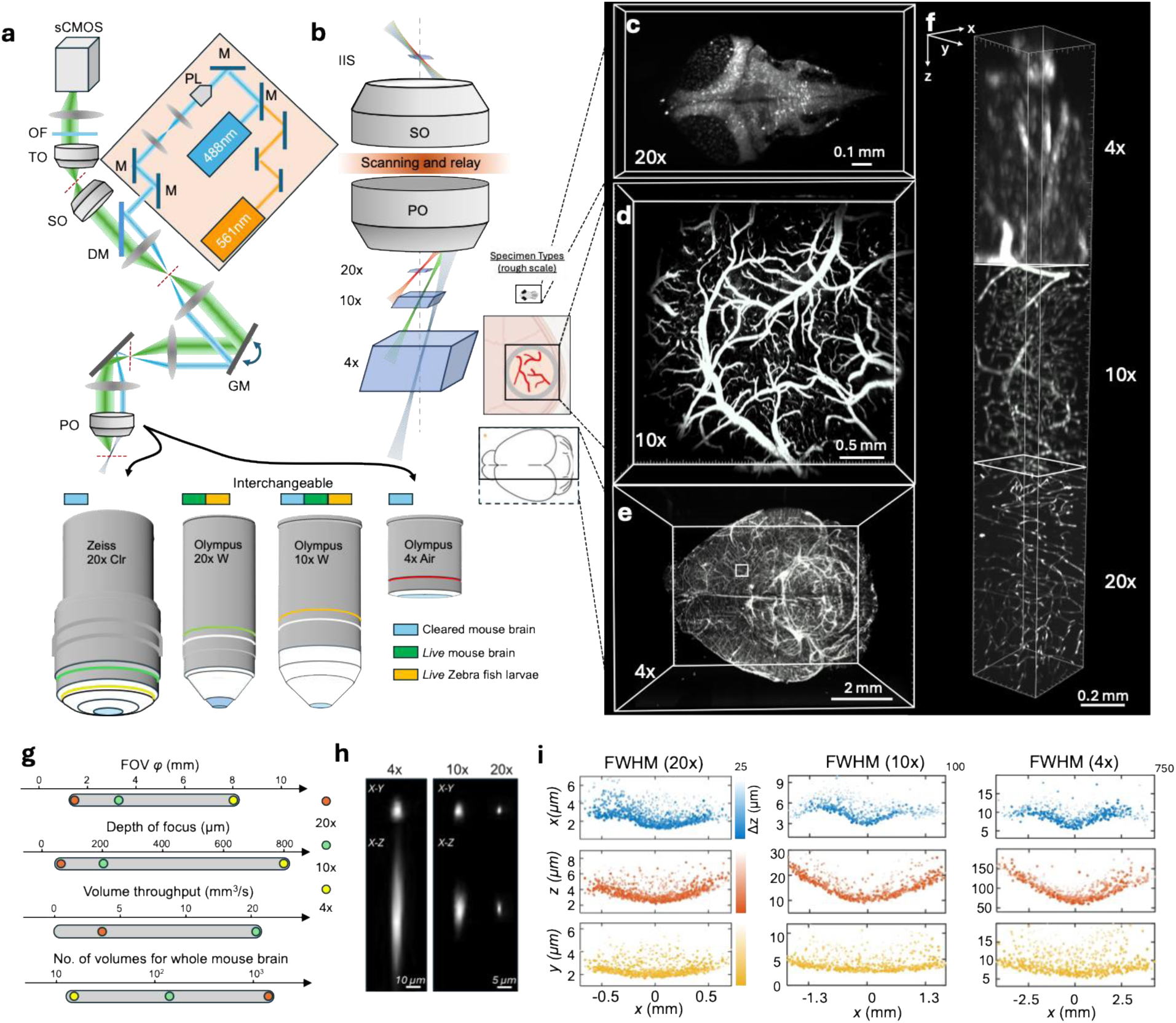
Self-adaptive mechanism of microscopic to macroscopic oblique plane microscopy. (a) The system layout schematic. The primary objective is switchable from 4× to 20×, with flexible immersive medium in air, water, and index matching liquid. M: mirror; PL: Powell lens; DM: dichroic mirror; GM: galvanometer mirror; PO: primary objective lens; SO: secondary objective lens; TO: tertiary objective lens; OF: optical filter. (b) The self-adaptive OPM optical path preserves a nearly invariant oblique angle at the IIS across multiple objectives, despite using different primary objectives with their distinct resolution and FOV specifications. Self-adaptive OPM was applied to image various sample types: (c) neural activity in a GCaMP expressing larval zebrafish brain imaged *in vivo* with the 20× objective, (d) vasculature of a FITC-conjugated dextran perfused mouse brain imaged *in vivo with the* 10× objective, and (e) GFP labeled endothelium of an optically cleared brain imaged *ex vivo* with the 4× objective. (f) A volumetric section of cleared mouse brain from (e) showing resolution improvement across the objectives. (g) The range of imaging parameters demonstrated in self-adaptive OPM. (h) Example point spread functions (PSFs) across the 4×, 10×, 20× objectives using fluorescent beads. Images were 10x interpolated for smooth display. (i) x, y, z resolution across the FOV and depth for the 4×, 10×, 20× objectives. Shading indicates the depth deviated from the focal plane.

OPM relies on a remote focusing method where the 3D object is re-imaged in the intermediate imaging space (IIS) and the tertiary imaging system refocuses the oblique light sheet onto a planar camera. The perfect imaging condition is conventionally obeyed, such that the remote image at IIS is free of optical aberration^9^. This condition requires that the lateral magnification from object space to IIS satisfies M_L_ = n_1_/n_2_, where n_1_ and n_2_ are the refractive index of the immersion media at the object and IIS respectively. Given the axial magnification is 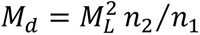, the magnification will be isotropic along all three dimensions as 𝑀_𝐿_ = 𝑀_𝑑_ = 𝑛_2_⁄𝑛_1_.

Instead of abiding perfect imaging conditions, we applied Abbe’s sine condition to unlock the flexibility of OPM. The oblique light sheet forms an angle *θ* with respect to the optical axis in object space, and its conjugate image forms an angle of ψ in the IIS (**Fig. 1b**). Abbe’s sine condition reveals the relation between the two angles by

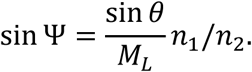

Assuming that the oblique angle is maximized by fully utilizing the NA of the primary objective (NA_1_), we can set sin 𝜃 𝑛_1_ = 𝑁𝐴_1_ and simplify as

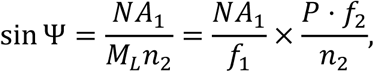

where *P* is the pupil magnification from the primary to secondary objectives, and 𝑓_1_ and 𝑓_2_ are the focal lengths of the primary and secondary objectives. This suggests that the conjugate angle of the oblique light sheet in IIS remains the same when the ratio of NA to the focal length of the primary objective is constant. This condition is easy to satisfy as long as the back aperture of the secondary objective is filled for the given primary objective. While the resolution is still fundamentally limited by the primary NA, Abbe’s sine condition allows switching between objectives for flexible choices of FOVs and resolutions.

We demonstrated four different objectives in this study, including Zeiss 20× clr (1.0 NA) specifically designed for optically cleared samples, Olympus 20× (1.0 NA) water immersion, Olympus 10× (effective 0.5 NA) water immersion, and Olympus 4× (effective 0.2 NA) air objective. For the 4× air objective, an immersion adaptor was designed to dip into liquid (**Supplemental Fig. S2**). Representative volume renderings of multiple resolutions and FOVs are shown in **Fig. 1c-1f**. The 20× magnification covers the entire larval zebrafish brain, including a portion of the spinal cord, as shown for a transgenic line in which neurons express a nuclear-localized GCaMP6 (**Fig. 1c**). The subcellular resolution (1.2×2×2.2 µm^3^ x-y-z) allows functional recording of single neuron activity across the entire brain of behaving larvae.

Mesoscopic volumetric *in vivo* imaging of hemodynamics in the mouse brain through a cranial window can be achieved using a 10× objective to image FITC-dextran perfused vasculature. A FOV of 3.2×3.2 mm^2^ and cellular resolution of 3×3×10 µm^3^ x-y-z is able to resolve circulating blood cells (**Fig. 1d**). Macroscopic scale *ex vivo* imaging was performed using the 4× objective (**Fig. 1e)** to image a whole mouse brain from a *Tie2-Cre:Ai9* reporter line expressing *tdTomato* in the endothelial cells. **Figure 1f** focuses on a single brain section (white box in **Fig. 1e)** to illustrate the comparison in 3D resolution across these three magnifications, all within the same system, by simply switching out the primary objective.

The same platform provides a FOV range spanning from 1.5 to 8 mm, and a depth of focus of 0.07 to 0.8 mm (**Fig. 1g**). At a ∼9,500 camera frame rate with rapid optical scanning, the volume throughput is up to 20.48 mm^3^/s, allowing 100 Hz imaging of a 3.2×0.32×0.2 mm^3^ local volume (and 3.2×3.2×0.2 mm^3^ global volume) of mouse brain (**Fig. 1g**). Using the 4× objective, the system facilitates rapid screening of optically cleared mouse brain volumes up to 15×20×7 mm^3^, with only 16 tiled volume acquisitions. The 20× objective, which is reserved for achieving the highest resolution on a local region of interest, would necessitate over 1,000 tiled volumes across the same coverage.

We characterized the 3D resolution for three different objective lenses using 0.2 µm fluorescent microspheres embedded in solidified agarose gel (**Supplemental Fig. S3**). Exemplary isolated bead images are shown in **Fig. 1h**. The resolution throughout the FOV along x for the three objectives is characterized in **Fig. 1i**, which shows the dependence of the FWHM resolution for the three dimensions with depth position encoded by color saturation. As we prioritized the FOV, the camera slightly under samples in the lateral *x* dimension so that the pixel pitch defines the resolution limit. The best resolution is achieved at the center of the FOV, at 1.2×2×2.2 µm^3^ for the 20×, 3×3×10 µm^3^ for the 10×, and 6×7×60 µm^3^ for the 4× objective. The axial magnification is a quadratic function of the lateral magnification, so the improvement in axial resolution is more significant with increasing magnification. The resolution worsens similarly toward the peripheral field across different objectives, even when satisfying the perfect imaging condition using the 20X primary objective. This indicates that aberrations from the relay optics and tertiary imaging systems are significant contributors to resolution worsening when pushing for a large FOV (**Supplemental Fig. S4**).

### 2. Mesoscopic volumetric imaging of cerebral vasculature

Cerebral blood perfusion provides metabolic support for brain activities. Imaging blood flow in the brain provides quantitative measures of perfusion, and opportunities to study the effects of neurological and vascular diseases. The high-speed OPM system with a mesoscopic FOV is an ideal tool to image *in vivo* dynamics of the mammalian brain which often requires coverage across an area of several millimeters. As a demonstration, we injected FITC-conjugated dextran into the retro-orbital sinus of an anesthetized mouse to label the vasculature and imaged through a cranial window.

**Figure 2a** shows a representative 3D rendering over a 3.2×3.2×0.2 mm^3^ mesoscopic volume imaged with the 10× objective. From the cross-sectional depth view (**Fig. 2b**), this microvasculature is discernible up to ∼200 µm below the surface of the cranial window. This depth range captures a physiologically critical zone that includes the surface vessel network and the initial penetrating arterioles that deliver blood into deeper tissue, combined with the microvascular plexus in cortical layer I^20^, offering insight into the metabolic and functional activity of the brain. Imaging at a 5 Hz volume rate with cellular resolution and color-encoded depth display (**Supplemental Video V1**) allows qualitative observation of blood flow dynamics at the capillary level. The intermittent signal along the capillaries is due to the absence of FITC-dextran signals from unlabeled circulating blood cells. To track the blood cell flow in capillaries over the entire FOV, we used a panoramic volumetric imaging protocol where a narrow 0.32 mm wide volume section, which sweeps the entire FOV in the slow scanning direction, is imaged at a 100 Hz volume rate (**Supplemental Video V2**). This approach maximizes the FOV uniquely accessible by mesoscopic OPM while still extending the 100 Hz volumetric imaging speed to the 10× objective to allow individual cell tracking.

**Fig. 2:**
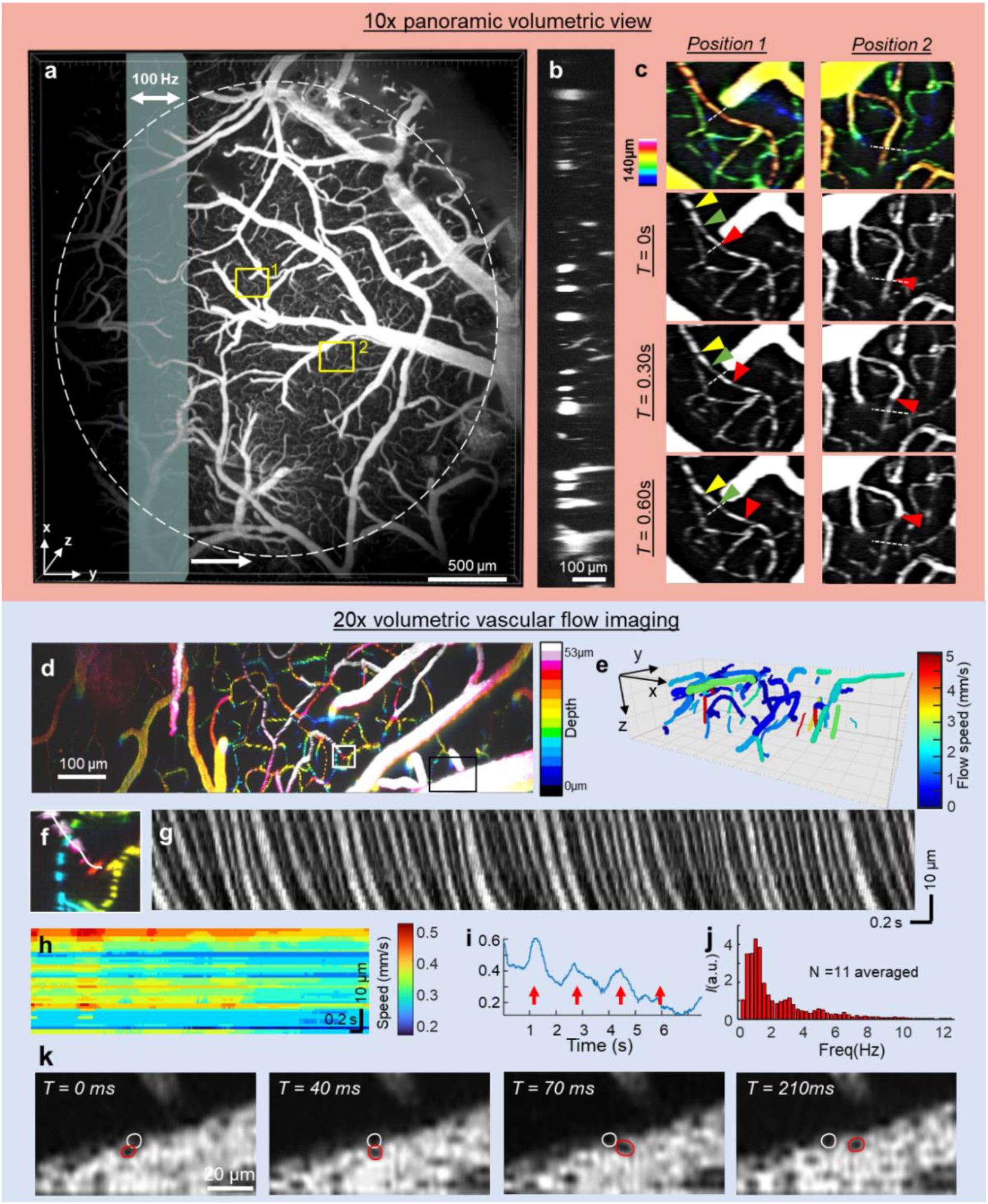
(a) 3D structural view of the mouse cortical cerebrovascular system captured with the 10× objective. The green bar indicates the local fast scanning volume section (100 Hz) sweeping the full FOV (right arrow). (b) Depth (x-z) cross section of the brain shown in (a). (c) Depth encoded maximum intensity projection of two small regions of interest from the yellow squares in panel (a) covering a depth of 140 μm below the surface of the cranial window. Snapshots at sequential timepoints show individual features (colored arrowheads) flowing through a capillary with differentiated flow characteristics and speeds. (d) Depth encoded maximum intensity projection acquired from the 20× objective. (e) 3D rendering of absolute flow speed of the local capillary network acquired with the 20x objective at 100 Hz volumetric rate. (f) Zoom-in of the white boxed ROI in panel (d) with a single vessel segment indicated (white line). (g) Kymograph collected from the capillary segment in panel (f) across 7 seconds in the collection. (h) Spatial and temporal dependence of blood flow speed from the kymograph in panel (g). (i) Changes in blood flow speed over time along a vessel segment showing a rhythmic pattern. (j) Frequency analysis of the speed profiles in panel (i) averaged from n=11 segments in panel (d) showing a dominant peak from 0.8-1.2 Hz. (k) A single depth position of the black boxed ROI in panel (d) at sequential time points shows two cells (red and white circles) as they roll past one another along the wall of a large vessel.

Capillary blood flow speeds are typically <5 mm/s, with cells traveling <50 µm between consecutive volumes at 100 Hz imaging rate, enabling deterministic tracking of individual cells^20,21^. In **Figure 2c** we sampled two ROIs from **Fig. 2a** (yellow boxes). The colormap encodes the depth of the capillaries, and the sequential snapshots show presumed blood cells flowing through the microvasculature network (**Fig. 2c**, arrowheads), permitting absolute quantification of speed. We switched to a 20× objective with better depth resolution and acquired time-lapse data at a 100 Hz volumetric rate over a 1.5×0.29×0.053 mm^3^ region, as shown in **Figure 2d** using color-encoded display of depth (**Supplemental Video V3**). Individual continuous capillary segments were identified and segmented within the network (representative zoomed segment shown in **Fig. 2f**). For each segment, the signal intensity along the vessel was extracted, creating a time-space diagram, or kymograph (**Fig. 2g**). As blood cells flow along the capillary segment over successive time points, the intensity contrast creates characteristic streak patterns in the kymograph, where the slope of these streaks is proportional to the projected flow velocity. The three-dimensional orientation of each capillary segment, determined from volumetric segmentation with the 20× objective, enabled calculation of the absolute flow speed along the capillary axis (**Fig. 2e)**. Here, we used the dominant slope provided by the Hough transform to estimate the average velocity for each vessel segment. This revealed vessel flow speeds up to ∼5 mm/sec, with most vessels displaying speeds around 2 mm/sec and slower, which aligns with the reported physiological range^21^. Penetrating vessels typically showed speeds at the higher end of this range.

The kymograph enables detailed spatiotemporal analysis of blood cell flow rates. This is achieved by using temporal correlation to calculate the travel delay between two locations on a capillary segment based on their intermittent flow patterns. This delay is then scaled by the segment’s length derived from the volumetric imaging data to determine the absolute speed. **Figure 2h** shows a representative spatiotemporally dependent capillary flow speed map from this analysis. Averaging the speed over the entire capillary segment (**Fig. 2i**) revealed rhythmic speed oscillations with peaks spaced approximately 1.3-1.5 seconds apart. This corresponds to a dominant frequency centered at ∼1 Hz in the speed profiles at each vessel position (**Fig. 2j**), a periodic pattern consistent with respiratory-coupled cerebrovascular dynamics. This coupling is driven by physical pressure gradients induced by breathing that propagate through the capillary network, and by shared chemical regulatory pathways between the respiratory an cerebrovascular systems^22,23^. This results in a ∼1 Hz oscillation that aligns precisely with the known breathing rate of anesthetized mice (55-65 breaths per minute)^24^. Our data strongly reveals the presence of this respiratory event in cerebrovascular speed measurements during *in vivo* imaging.

Large arterioles and venules present a distinct challenge. Blood cell velocities in these vessels often exceed the tracking capacity of 100 Hz volumetric imaging. However, laminar flow in larger vessels creates a substantial velocity gradient near the vessel wall, where 100 Hz 3D imaging remains capable of resolving deterministic cell movements. **Figure 2k** illustrates such an example, showing a single depth section at sequential time points where two individual blood cells, presumably immune cells undergoing margination, roll past one another along the wall of a large vessel before being re-entrained into the faster central flow stream. This capability to simultaneously observe distinct flow regimes, from tracked cell motion in capillaries to resolving margination dynamics in larger conduit vessels, demonstrates diversity of vascular flow phenomena accessible through mesoscopic OPM imaging.

### 3. Volumetric calcium imaging on behaving zebrafish larvae

To further showcase the capabilities and versatility of this OPM system, we recorded the dynamics of a genetically encoded fluorescent calcium sensor (GCaMP) of behaving larval zebrafish, by both in toto whole body imaging (10x) and brain-wide single neuron imaging (20x). Larvae were exposed to a grating pattern moving in the tail-to-head direction to trigger the Optomotor Response (OMR), an innate behavior in which the experience of backwards self-motion within a stationary environment evokes forward swim bouts to oppose this perceived motion^25,26,27^. OMR has been studied in many species, and has been observed as early as 5 days post fertilization (dpf) in zebrafish^28^. We used a custom designed experimental setup similar to previously described methods^29^ to simultaneously present the OMR stimulus, capture tail movements, and image calcium dynamics (**Fig. 3a)**. A moving grating stimulus was displayed on an opaque screen below the fish to depict whole visual field movement, an infrared camera was aligned below the fish to record the tail, and the OPM objective above the fish captured calcium activity.

**Fig. 3:**
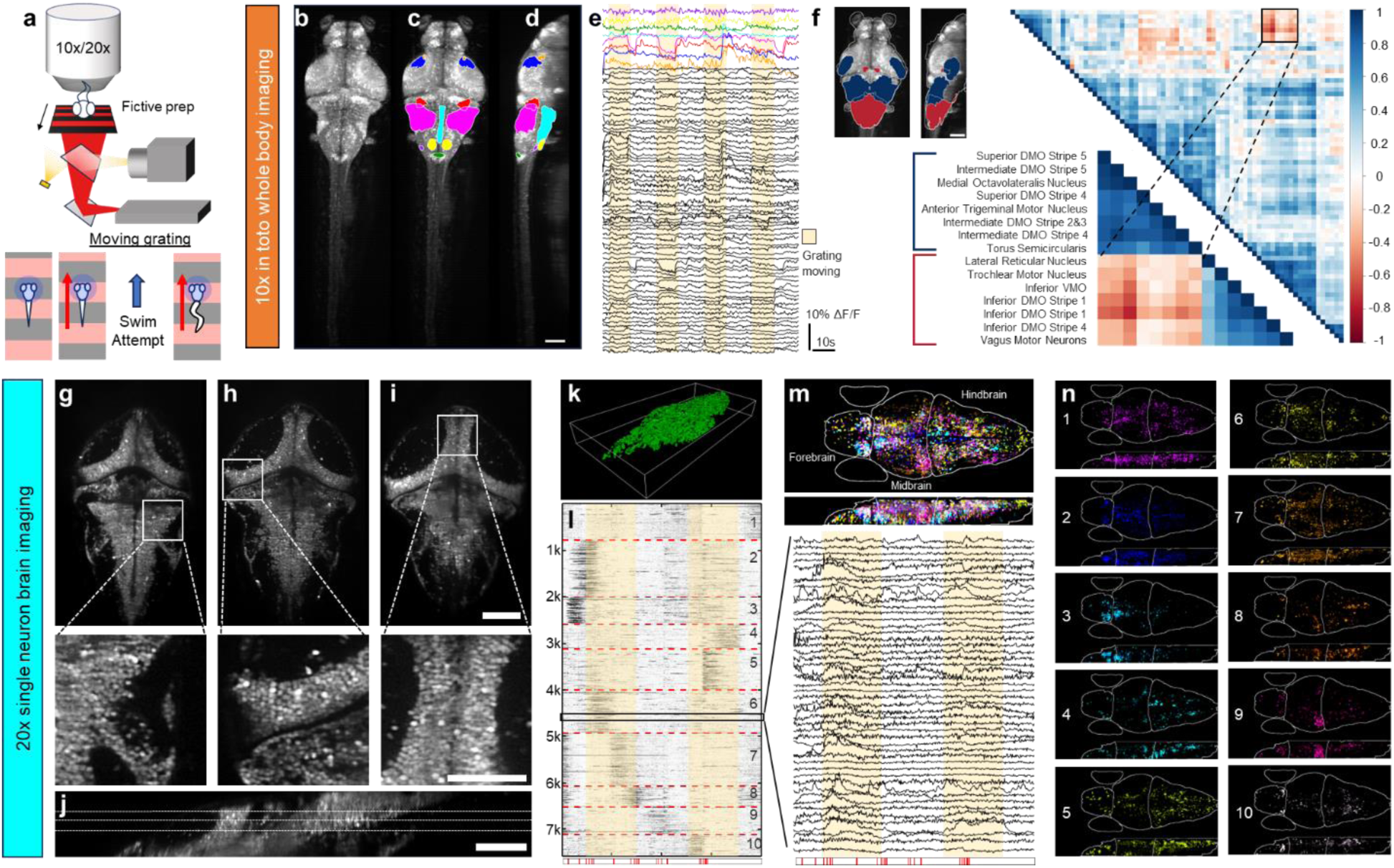
Zebrafish Calcium Imaging During the Optomotor Response. (a) Simultaneous OMR stimulus projection and whole body recording in a fictive prep experimental setup for OPM imaging. A stimulus grating is reflected by a mirror onto an opaque surface below the fish after passing through a cold mirror. The cold mirror reflects an infrared light source back to a camera to capture tail movements. The OPM objective is positioned above the fish. (b) Maximum intensity projection of calcium signals from a whole body volume scan of a zebrafish larvae expressing GCaMP7b in neurons throughout the nervous system imaged with the 10× objective. Discrete anatomical regions defined by the MZB brain atlas overlaid on (b) viewed from above (c) and from the side (d). Scale bar = 100 µm. (e) Average intensity traces from selected regions color-coded per (c) and (d). Black traces are from areas outside of the regions delineated in panels (c-d). (f) Heatmap of cross correlation between discrete regions, inset shows specific regions with correlated and anticorrelated activity patterns and their anatomical locations within the brain. DMO, Dorsal Medulla Oblongata, VMO, Ventral Medulla Oblongata. (g-i) Single slices from a volume of a larval zebrafish expressing GCaMP6f in the nuclei of almost all neurons imaged using the 20× objective with insets showing single neurons. Scale bar = 100 µm for full images and 25 µm for insets. (j) Single slice view of the image volume viewed from the side, dotted lines indicate planes shown in panels (g-i). (k) 3D rendering of segmentation mask of individual neurons. (l) Raster plot of individual neuron traces organized by cluster, zoomed inset shows individual traces, red heatmap below indicates the timing of tail movements. (m) Color-coded segmentation mask, each color represents a specific neural response cluster, color intensity indicates relative contribution of that neuron to the cluster activity pattern. Dorsal view top, side view bottom. (n) Individual clusters from (m) separated to highlight distinct anatomical regions represented in each cluster.

The mesoscopic FOV offered by the 10× objective enabled whole-body imaging of 7 dpf larvae with an approximate 2×0.4×0.35 mm^3^ volume containing the entire brain and spinal cord of larvae expressing GCaMP7b in neurons (**Fig. 3b-3c**). Notably, the depth of focus extends the entire thickness of the larval brain (**Fig. 3d**). Volumes were captured at 5 Hz for a total duration of 80 seconds while the OMR grating stimulus alternated between moving and still. Volumetric data was then registered to the zebrafish brain atlas MapZeBrain (MZB) and GCaMP traces were extracted from 74 non-overlapping anatomical regions (**Fig. 3c-3e**).

The neural activity from different regions across the entire brain were analyzed and quantified through a correlation matrix (**Fig. 3f**). Throughout the OMR assay, one area spanning the posterior portion of the midbrain/optic tectum (OT) and anterior hindbrain had activity patterns highly anticorrelated with another region covering the posterior hindbrain (**Fig. 3f**, boxed inset). Although the OT is not required for the OMR^30^, we observed intensity spikes in superficial OT layers and extratectal arborization fields during stimulus motion (blue and orange traces respectively, **Fig. 3e**). These regions are known to be targeted by the axons of retinal ganglion cells (RGCs) involved in transmitting visual field motion information^31–34^ and are therefore expected to be activated by the OMR stimulus. In the hindbrain, subregions within the inferior Dorsal Medulla Oblongata (DMO) showed drastic drops in activity correlated with stimulus motion onset or offset (magenta and red traces, **Fig. 3e**). Neurons in these regions have been shown to be responsible for coordinating sensory and motor information and were likely activated by the fish attempting to swim against the motion of the stimulus^35^.

While averaged GCaMP traces allow regionalized whole brain analyses, the small tightly packed neurons in the larval zebrafish brain (typically 5-7 µm in diameter) require sub-cellular resolution to dissect brain activity at the single neuron level. To do so, we switched to the 20× objective to perform calcium imaging of nuclear localized pan-neuronal GCaMP6f during OMR stimulus presentation. A 940×430×100 µm^3^ region covering the brain with single cell resolution (**Fig. 3g-3j**) was imaged at a 5 Hz volume rate (**Supplemental Video V4)**. The near isotropic 3D sampling density deterministically located single nuclei within the depth of focus. We acquired a static structure volume image and applied 3D watershed segmentation to isolate >7,500 single neurons (**Fig. 3k**). We subsequently applied the resulting 3D masks to the registered time-lapse volumetric data to extract individual neuron GCaMP traces (**Fig. 3l**) over the 80 second period of the OMR assay.

Individual traces were grouped into 10 clusters using Non-Negative Matrix Factorization as previously described^35^. Each resulting cluster contained approximately 500-1000 neurons and neurons within each cluster showed one or two short periods (5-10 s) of high activity during the 80 second assay (**Fig. 3l**). Representative traces from 100 neurons showed rich and diverse activity patterns that were diluted by region-averaged whole-brain analysis from the 10× objective (**Fig. 3l**, inset). Mapping the clusters back to the original mask revealed distinct anatomical locations for many of the neurons found within a single cluster (**Fig. 3m-3n**). We then correlated individual clusters to both the stimulus pattern and fish tail movements recorded by the infrared camera (**Supplemental Video V5**). Cluster 2 neurons showed high levels of activity that coincided with routine, small swim bouts observed at the beginning of the assay. The cells of this cluster are highly represented in the nucleus of the medial longitudinal fasciculus (nMLF) and anterior hindbrain (aHB), both of which have been shown to be active during visually driven forward swims^36^. A single struggle-like swim bout followed by a lack of responsiveness was observed in the second half of the assay, and clusters 10 and 5 had activity correlated to before and after this struggle respectively. Many of the cluster 10 neurons mapped to the locus coeruleus (LC) or noradrenergic cluster of the Medulla Oblongata (NE-MO), both of which have been shown to increase activity prior to a switch to passivity. Although cluster 5 neurons are spread across the entire brain, there is a strong representation of the cluster in the lateral MO (L-MO) which likely corresponds to the inhibitory GABAergic neurons that suppress motor outputs during this passive state^35^. Additionally, cluster 6 shows activity most highly correlated with the onset of stimulus motion. Many of these neurons are located in the pretectum, which is known to be involved in the initial perception of visual stimuli such as those presented during the OMR assay. Clusters 4 and 8 showed activity during the later stages of stimulus movement and appear to be localized to regions of the hindbrain containing the ventromedial spinal projection neurons (vSPNs) responsible for initiating motor responses for forward swims associated with OMR^36,37^. Clusters 1 and 7 showed the lowest changes in activity throughout the assay and are also relatively evenly distributed across the brain, likely corresponding to neurons that are spontaneously firing and not associated with OMR. Overall, agreement between this data and previous studies of the OMR in zebrafish establishes OPM as a reliable and flexible system for performing diverse whole-brain analyses in behaving larval zebrafish.

### 4. Multi-scale volumetric imaging on optically cleared and mildly expanded mouse brain

Tissue clearing has become a powerful tool in biological studies because it homogenizes refractive indices and suppresses scattering, enabling high-contrast, high-resolution imaging of whole tissues and organisms^38,39^. Tissue expansion further improves effective resolution by isotropically enlarging the specimen and separating nearby subcellular structures so that features originally below the optical diffraction limit became physically larger and resolvable^40^. Processing the massive volume of data generated by subcellular-resolution imaging poses a data efficiency challenge. Typically, features within a sample have different imaging demands, such as high-resolution imaging for subcellular features in sparse cells or large volume imaging for connectivity of long-range projection neurons. The flexibility to switch the volume field and resolution in the self-adaptive OPM enables stratified imaging from macroscopic coarse screening to local microscopic high-resolution imaging (**Fig. 4**).

**Fig. 4:**
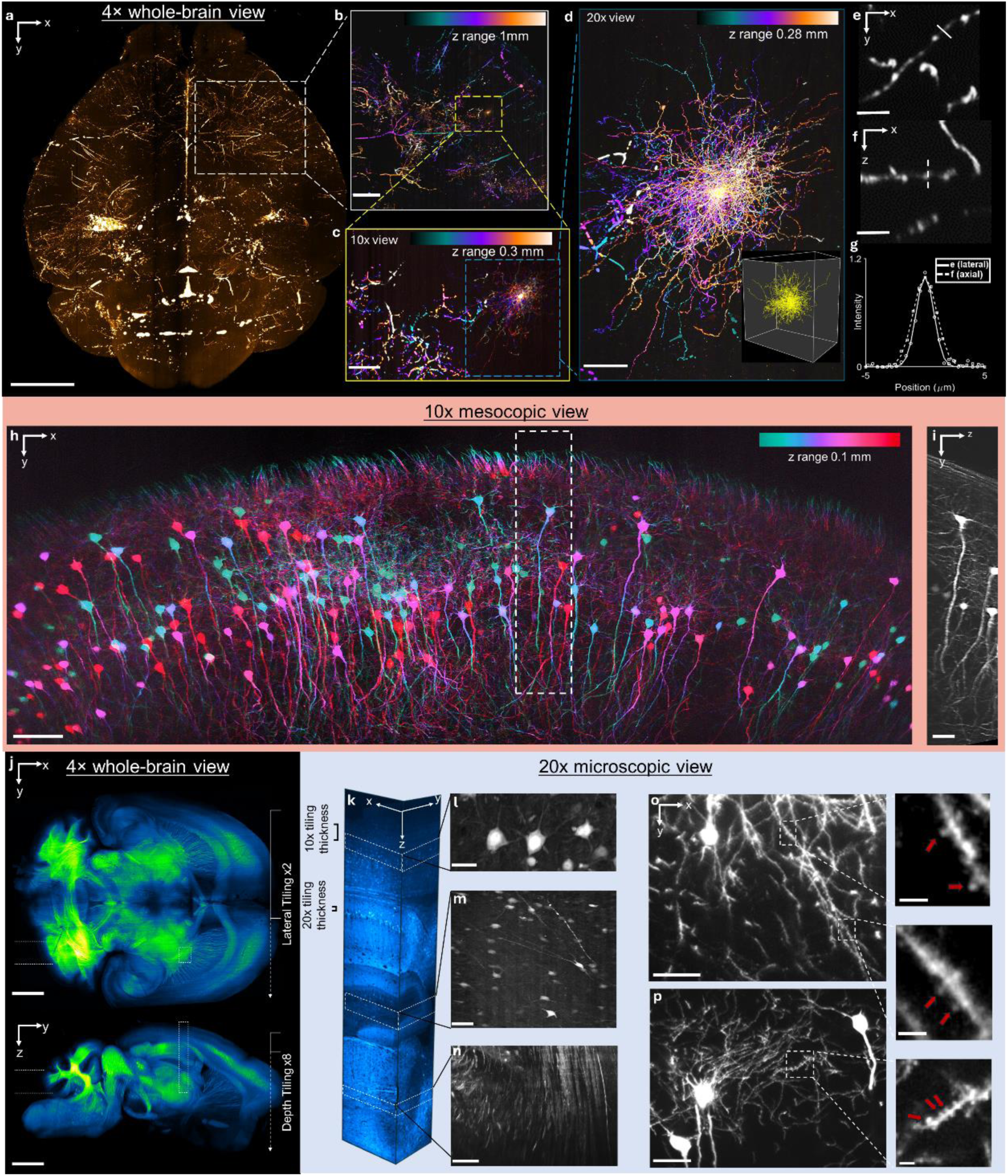
Multiscale Volumetric Imaging of Cleared Mouse Brains. (a) 4× x-y overview of a sparsely labeled *tdTomato*-expressing adult mouse brain optically (RI = 1.496). Maximum intensity projection of the central 4 mm depth volume, stitched from 8 lateral × 2 vertical tiles with 10% overlap. Scale bar = 2 mm. (b) Depth color-coded maximum x-y projection from the boxed region in right posterior dorsolateral parietal cortex (z depth 1 mm), highlighting a multipolar neuron and subcortical structures. Scale bar = 500 µm. (c) Boxed region of interest imaged at 10× magnification, depth color-coded maximum x-y projection over 0.3 mm z depth. Local arbors and collaterals become resolvable while maintaining continuity with mesoscale tracts. Scale bar = 200 µm. (d) 20× magnification x-y view of a single local inhibitory interneuron in layer 2/3 of the parietal cortex. Depth color-coded maximum projection of a 0.28 mm z volume, assembled from 5 lateral tiles; inset shows the 3D surface rendering of the entire cell. Scale bar = 100 µm. (e-f) Magnified dendritic segments from the neuron shown in (d) from the x-y view (e) and x-z view (f). Scale bar = 20 µm. (g) Plot of intensity profiles along the solid white line in (e) and dashed white line in (f). Measured resolution from FWHM of the Gaussian fit is 1.8 µm lateral and 2.2 µm axial. (h) 10× depth color-coded maximum projection from a mildly expanded Thy1-GFP brain, revealing putative CA1 pyramidal neurons and local processes (z range = 0.1 mm). Scale bar = 100 µm. (i) Orthogonal view (y–z) from the boxed region in (h). Scale bar = 40 µm. (j) 4× mesoscopic maximum intensity projections (dorsal and sagittal) of a Thy1-EGFP cleared brain (RI=1.52). Total 8 lateral × 2 vertical tiles with 0.75 mm lateral step size and 10% overlap. Scale bar = 2 mm. (k) 20× objective acquisition of a columnar section from the boxed section in (j) ranging from the right superficial cortical area to thalamus acquired from 68 lateral tiles covering a depth of 3.5 mm along z. (l-n) Sub-regions from the 20× acquisition in panel (k) shown in x-y view highlighting (l) cortical neuron cell bodies and proximal dendrites (scale bar = 25 µm), (m) long axonal projections crossing corpus callosum to hippocampal pathways (scale bar = 100 µm), and (n) thalamic fibers (scale bar = 100 µm). (o-p) 20× imaging from the brain in (h-i) resolving dendritic trees and higher order branches. Detail insets (right) highlight spine-like protrusions and spine heads (red arrows). Scale bars = 35 µm (o–p) and 4 µm (insets).

**Figures 4a-4d** show an example of a sparsely labeled, *tdTomato* expressing adult brain cleared to a refractive index (RI) of 1.496 with increasingly magnified structural detail. We first acquired a 4× overview of the entire brain using 16 tiles to cover a volume of 12×15.5×6.5 mm^3^ (x-y-z). *En face* maximum intensity projection across the central 4 mm depth section is shown in **Figure 4a**. A more detailed view of this image (**Fig. 4b**) focused on the right posterior dorsolateral parietal cortex (depth-color coding across z range of 1 mm) revealed a bright neuron of interest, likely a radially symmetric multipolar neuron. To further examine this neuron, we reimaged the identified target with the 10× objective (3×4×0.3 mm^3^, x-y-z), which preserved continuity with mesoscale tracts while resolving local arbors and collaterals (**Fig. 4c**). Combining the 4× mesoscale image with the detailed 10× structural image confirmed the presence of a local inhibitory interneuron in layer 2/3 of the parietal cortex.

To capture even finer details of this neuron, we switched to the 20× objective and imaged across a FOV of 1.5×2×0.75 mm^3^ (x-y-z) split into 14 lateral tiles. The depth color-coded projection of a 0.28 mm depth volume section assembled from five lateral tiles is shown in **Figure 4d**. The full-cell 3D surface rendering (**Fig. 4d**, inset) assembled from all 14 tiles confirmed the complete morphology. Line profiles through dendritic segments from the 20× image (**Fig. 4e–f**) yielded a FWHM (**Fig. 4g**) of 1.8 µm laterally and 2.2 µm axially, in agreement with the system’s resolution characterization. This sequential workflow and multiscale volumetric imaging flexibility enabled rapid morphological examination of a single cell in a long-range circuit, linking local structure to macroscale organization.

We next imaged a *Thy1-CreERT2-*EYFP (SLICK-H) transgenic mouse brain cleared to RI of 1.52. A quick 4× overview scan achieved full coverage of the entire brain (10.5×12.5×6 mm^3^, x-y-z) shown by the maximum intensity projection in the dorsal (x-y) view and sagittal (x-z) view (**Fig. 4j**). A registered 20× column (acquired from 68 tiles) spanning from the right superficial cortex to the thalamus covered 3.5 mm in z (**Fig. 4k**). Subfields illustrated cell bodies with proximal dendrites (**Fig. 4l**), long axonal projections crossing the corpus callosum toward hippocampal pathways (**Fig. 4m**), and thalamic fibers (**Fig. 4n**). An additional acquisition centered on the cerebellum reproduced the cerebellum trilaminar architecture (molecular layer, Purkinje cell layer with planar dendritic fans, and granule cell layer)^41^ and delineated the cerebellar peduncles that bridge the cerebellum to the brainstem (**Supplemental Fig. S6**). This cross-depth readout provides quantitative biomarkers relevant to cerebellar pathology, such as loss of dendritic planarity in Purkinje cells and peduncular fiber rarefaction, which are common in neurodegenerative disorders including multiple system atrophy cerebellar type and spinocerebellar ataxia^42–46^.

A second GFP expressing Thy1 brain was optically cleared with Cubic R+ RI matching solution to produce 1.5× linear expansion and 3.5× volume expansion of cellular and neuropil structures^47^. The cleared brain was cut sagitally at the midline to increase access to deep brain regions like the cornu ammonis 1 (CA1) and the dentate gyrus (DG). Under these conditions, imaging first with the 10× objective resolved CA1 pyramidal neurons and local processes within the hippocampus (**Fig. 4h-i**). Increasing the magnification to 20× in a different CA1 region resolved dendritic trees with higher order branches and spinelike protrusions **(Fig. 4o-p**) and revealed individual dendritic spines with identifiable spine heads. The modest clearing-induced expansion effectively scaled optical resolution, enabling direct observation of isolated spines without requiring dedicated expansion microscopy.

## Discussion

We present a different design principle in OPM that allows self-adaptation to desired FOV, resolution, and imaging volumes. Like the operation of a conventional microscope, OPM allows for switching of the primary objective at will to zoom at different magnifications and fit specific application needs. Compared to other state-of-the art systems, self-adaptive OPM achieves the highest volumetric throughput across all FOV capabilities (**Supplemental Fig. S7 and Supplemental Table 1**). The FOV diameter ranges from 1.5 to 8 mm. The corresponding lateral resolution spans from 1.2 to 6 µm, depth resolution from 2 to 60 µm, and depth of view from ∼70 to 700 µm, covering microscopic to macroscopic volumes.

The highest resolution is capable of brain-wide volumetric calcium imaging in behaving larval zebrafish. The 20× objective satisfied the perfect imaging condition and has near isotropic resolution in 3D, allowing single neuron segmentation and deterministic extraction of calcium traces and circuitry analysis on behaving animals. Mesoscopic volumetric imaging with the 10× objective provides cellular resolution imaging of the mouse brain, accessing a ∼3×3 mm^2^ FOV. We demonstrated a 5 Hz volume rate over the whole FOV, and a 100 Hz volumetric imaging using a panoramic imaging method. Given a typical neuron soma size of 20 µm, the cellular resolution (3×3×10 µm^3^ in x-y-z) and mesoscopic FOV enable interrogation of large-scale dynamics in mouse cortex rooted in single-cell activity. Extending to macroscopic volumetric imaging, we showcased the capacity to rapidly cover the entire optically cleared mouse brain through mechanical scanning. The rapid and coarse volume survey can first identify ROIs and then high-resolution data can be obtained with the 20× objective specifically designed for cleared samples. Both multi-scale flexibility and high-speed dynamic capability make this imaging method well-suited for a wide range of biological applications in complex 3D whole organisms, such as other model systems like C. elegans and Drosophila, as well as in zebrafish and mouse brain as demonstrated here.

We have previously shown that the aberration in IIS is independent of the primary objectives, particularly with lower NA options, when employing remote imaging^12,48^. Building on this foundation, our design prioritized the FOV and achieved a field number (FN) of 30, exceeding the designed specification of FN 22. The 3D resolution characterization revealed that the peripheral aberration pattern was remarkably similar across all primary objectives regardless of adherence to the perfect imaging condition. Thus, when pushing for large FOV, the aberration introduced by relay optics emerged as the dominant factor limiting the achievable resolution. With off-the-shelf optics, we opted to tolerate the aberration. For the demonstrated imaging of intact multicellular organisms, the resolution appears capable of resolving individual cells, or even subcellular structures in optically cleared samples. The focus of this work is to demonstrate self-adaption of OPM under Abbe’s sine principle, which alleviates the constraints imposed by the perfect imaging condition. By leveraging advanced approaches to improve resolution, either by computation or physical expansion, the optical design becomes more forgiving and opens the door to further push imaging volume or imaging speed.

There are several improvements that can be immediately implemented in the remote imaging system, such as using refractive index mismatching tertiary objectives to more efficiently collect light and improve resolution^11^, simultaneous two-color imaging for multi-channel excitation^7^, or introducing a variable tube lens beyond the tertiary objective to adjust the zoom factor^49^. We also propose a more careful design of the relay optics and tube lens to improve the overall imaging performance. Two photon scanning light sheet excitation can increase the penetration depth into scattering samples. Remote axial focusing can be introduced to correct sample curvature^50^. The simplicity of self-adaptation in OPM opens broad possibilities for imaging 3D multi-cellular whole organisms.

## Methods

### 1. System Architecture and Optical Configuration

#### 1.1. Imaging System

Light sheet illumination was generated by passing the collimated laser output from the OBIS laser (488 nm) through a Powell lens (60° fan angle, LGL160, Thorlabs). The resulting illumination was reshaped by a 3:10 telescope (Thorlabs AC254-030-A, f = 30 mm; Thorlabs AC254-100-AB, f = 100 mm). A second OBIS laser line (561 nm) was co-aligned with this path using a pair of kinematic mirrors. These illumination components were installed on a translatable optical breadboard to fine-tune the light sheet illumination offset position for each objective. The illumination laser path was then steered by an additional kinematic mirror pair for fine alignment. A multi-band dichroic mirror (Di01-R405/488/561/635/800-t3-25x36, IDEX Semrock) reflected the illumination into the main optical path. Two 1:1 relay telescopes mapped the Fourier plane of the light sheet onto the scanning galvanometer mirror with a 20 mm diameter optical clearance, and then to the back aperture of the primary objective. Each of the four lenses in these two relay steps consisted of a lens doublet (2x Edmundoptics 49392, 50 mm diameter, f = 200 mm, VIS-NIR coated achromat) with a combined focal length of 100 mm. A mirror directed the beam downward, creating an upright microscope configuration. The secondary and tertiary objectives were 20×, NA=1.0 water immersion lenses (Olympus XLUMPLFLN20×W). A water bath was maintained between these objectives using an elastic rubber sheet seal. The tertiary imaging system was comprised of the tertiary objective lens, emission filter, tube lens, and high-speed sCMOS camera (Kinetix22, Teledyne Photometrics). The tube lens compound assembly consisted of a doublet (Thorlabs ACT508-300-A, 50 mm diameter, f=300 mm) combined with a tube lens (TTL200MP, f = 200mm), yielding an effective focal length of 120 mm.

The magnification for the tertiary imaging system was 13.3×. The entire detection system was mounted on a 3-axis translation stage to adjust the translational position of the detector, and a custom rotational platform to adjust the detection angle about the fixed IIS positional axis. A detailed sketch of the optical layout is included in **Supplemental Fig. S1** with all parts listed in **Supplemental Table 2**.

#### 1.2. System control

Acquisition control was managed via custom LabVIEW software. A TTL signal, synchronized to the frame exposure onset, was output by the camera and served as the external clock on a NI DAQ board analog output channel (USB 6363). A sawtooth voltage waveform, pre-programed with the desired step numbers and duty cycle, drove the galvanometer for optical scanning. All waveforms were automatically synchronized with the camera frames through the external clock trigger.

### 2. Mouse Cerebrovascular Imaging

#### 2.1. Mouse Surgical Preparation

Mice underwent cranial window surgery for *in vivo* cerebrovascular imaging. A 4×4mm^2^ circular section was carefully removed from the mouse skull without perforating the dural tissue layer. A small drop (10 µL) of 1% low melting point agarose was placed directly on the brain within the section of skull that was removed. Before the agarose set, a 5 mm diameter cover slip was placed over the craniotomy and glued in place, ensuring that the cover slip was flush with the skull on all sides. A headpost was mounted to the contralateral side of the skull, and the entire surgical site was sealed with Metabond cement. Imaging was performed on the same day as window implantation to minimize the impact of inflammation on image quality. To prepare for imaging, animals were held under isoflurane anesthetic and received a 100 µL injection of 5% Fluorescein Isothiocyanate Dextran (FITC-dextran, Sigma Aldrich, MW 2,000,000) into the retro-orbital sinus. The animal was head restrained on a 3-axis translation stage and the position of the brain was adjusted to co-align it with the sample plane of the microscopy system.

#### 2.2. Dynamic Imaging Acquisition

Structural reference images were first acquired at low laser power (488 nm, 2 mW) and an extended exposure (5 ms) to serve as an alignment target for subsequent dynamic imaging and post hoc image registration. For the 10× objective, structural imaging covered a 3.1 mm range (2.2 V) along the y-scanning dimension in 512 frames (6 µm step size). The 20× objective scanned 0.29 mm (0.4 V) in 512 frames (0.56 µm step size). The NI DAQ board synchronized the camera triggering and galvanometer step position for each slow scan step position by incrementally scanning across the full y-scanning range. The camera was operated in the “Edge Trigger” mode so that the camera would wait a TTL signal for each step position to begin a frame exposure.

After the alignment was optimized from the structural information, dynamic images were acquired. For the 10× objective, structural resolution was maintained but the exposure was reduced to 0.37 ms and laser power was increased to 20 mW to achieve 5 Hz volumetric imaging. For the 20× objective, the sampling density was reduced to 3 µm spacing (95 frames) and exposure dropped to 0.1 ms with 20 mW laser power to achieve 100 Hz volumetric imaging. Here, the camera operated in “Trigger First” mode to maximize the frame rate. This mode requires only a single trigger to start the acquisition, which then runs continuously until all frames are collected. The start of the first camera row exposure delivered a TTL signal to the NI DAQ board to trigger the galvanometer to begin its scan sequence, which ran in a loop during the acquisition. Extended acquisition parameters are included in **Supplemental Table 3.**

#### 2.3. Panoramic Volumetric Scanning

To simultaneously achieve high volumetric rate imaging of capillary flow while maintaining full-field coverage of the 10× objective, we implemented a rolling-scan acquisition strategy that leverages the high spatial resolution of the 10× objective to resolve single-cell dynamics while preserving network-wide context. Sequential imaging volumes were acquired with incrementally offset starting positions, enabling high-speed local scanning (100 Hz) across the entire cranial window at a slower effective global rate. To achieve this, a second analog output from USB-6363 produced a TTL square waveform for every sawtooth cycle and served as the external clock for a secondary NI DAQ board (USB-6002). A ramping voltage signal was programmed to sweep the volumetric imaging across the entire FOV. The ramping and sawtooth waveform were combined by an additive op amp circuit before being fed to the galvanometer driver board.

Each local volume covered 0.32 mm (0.23 V) with 3.4 µm sampling density (95 frames) at 100 Hz volumetric rate (0.1 ms exposure per plane). Adjacent volumes overlapped by 99.5%, requiring 1,540 acquisitions to span the full y-scan range and yielding a total acquisition time of 15.4 seconds per complete volume. The acquisition speed could be increased by reducing the overlap between successive scans; for example, reducing to 99% overlap lowered the acquisition time to 7.7 seconds (770 local scans). To create the final output video, the background was populated from the structural image. Each active scanning region was y-translation shifted to align with the background. To better visualize the signal in the active region, a contrast enhancement transformation was applied to each active window (Matlab: adapthisteq with ‘NumTiles’=16x16 and ‘ClipLimit’=0.01).

#### 2.4. Volumetric Mapping of Flow Speed

To analyze the neovascular flow dynamics, first a pseudo-structural image was generated from the average of the first fifty dynamic volumes. A 3D Gaussian filter was applied to remove background noise and enhance the vessel contrast. A manual threshold was applied to create a mask for segmentation. Residual floating pixels and gaps were removed (MATLAB “bwareopen” and “imclose” with sphere structuring element respectively). This mask was skeletonized (MATLAB “bwskel”) and segmented into starting points and branching points to identify discrete vessels (MATLAB “bwmorph3” and “bwlabeln”). The diameter of each vessel was estimated (MATLAB “bwdist”) to create a diameter map corresponding to each pixel in the structural image. Any vessels narrower than the threshold were discarded. The output segmentation was manually reviewed and additional branchpoints were added to ensure labels were correctly assigned and matched the structural “ground truth” acquisition.

The segmentation coordinates were unidirectionally sorted using a 3D distance minimization approach. The endpoint provided by bwmorph3 was used as the starting point. The next closest point was identified from the minimum distance between all other points. The direction was calculated between subsequent points to eliminate directionality reversal along the sorted coordinates. This process was repeated until all points in the segmentation were sorted. The total length of the vessel segment was estimated using the sum of the distance between subsequent points. The x and y coordinates from the vessel segmentation were then used to define a kymograph by capturing the intensity timeseries from the 2D maximum intensity projection of the volumetric timeseries at each vessel position. In the kymograph, the spatial extent of the segment is represented along the y-axis and time is represented along the x-axis. The angle of the resulting intensity streaks along the kymograph is used to find the flow speed of particles moving along the vessel. Each kymograph was visually inspected to confirm unidirectionality in the flow profile. If there was an abrupt change in the flow angle, this indicated a missing vessel branch from the automatic segmentation, and the segmentation and kymograph were re-generated with the new branch point.

Each kymograph was pre-processed with standard Gaussian denoising, edge enhancement through Sobel filtering, and contrast enhancement by stretching to the 2^nd^ and 98^th^ percentiles. The Canny edge detection method created a binary edge image to input into the Hough transform to find the dominant angle (θ) and its position (ρ). This angle was converted to velocity by taking the sine of the dominant angle and scaling by the pixel length and the time per frame from the acquisition settings. The pixel length from the microscope calibration was scaled by the 3D segment length to consider the distance traveled along all directions, including depth. The average velocity for each segment was mapped back to the 3D segmentation.

#### 2.5. Spatiotemporal Analysis of Vessel Segments

To characterize spatiotemporal flow fluctuations within vessel segments, the strongest segments were manually selected from the x-y projected volumetric timeseries.

Kymographs were generated by defining equally spaced coordinates along the segment centerline. Mean intensity profiles were calculated along the tangent direction at each position to account for frame-to-frame misalignment. Flow dynamics were quantified using the cross-correlation between two spatially offset locations along the segment.

The temporal window size was set using the dominant flow angle from the Hough transform to estimate the time for cells to traverse the vessel segment. The window size was at least twice this transit time to ensure sufficient data for correlation analysis. The lag corresponding to the dominant peak in the cross-correlation was recorded to calculate time delay. The speed was then calculated by dividing the distance between two locations by the time delay. This window was shifted incrementally across the kymograph to visualize flow changes at each spatiotemporal position. For frequency analysis of the speed change over time, the temporal speed profile was fitted by 3^rd^ order polynomial which was then subtracted. The resulting profile was smoothed and then Fourier transformed to reveal dominant frequency from periodic flow patterns at capillary segments.

### 3. Larval Zebrafish Neural Calcium Imaging

#### 3.1. Zebrafish Care and Preparation

Transgenic larvae expressing the genetically encoded calcium indicator GCaMP in the cytosol (Tg(elavl3:jGCaMP7b), **Fig. 3a-f**) or nucleus (Tg(elavl3:H2B-GCaMP6f), **Fig. 3g-o**) of most neurons were used for zebrafish imaging experiments. All larvae were in the Casper pigmentation background^51^ to lack melanophores and iridophores and remain optically transparent. Fish were raised following standard procedures (28°C in E3 media, light cycle conditions of 14 h light and 10 h dark, Paramecia feed starting from 3 dpf) and imaged at 7 days post fertilization (dpf). For imaging, larvae were transferred to heated 2% low-melting agarose and mounted in the center of a Petri dish with the dorsal side facing up. For whole nervous system experiments (10×), the full fish was embedded in agarose to keep the tail immobilized during the recording period. For whole brain experiments (20×), a tail-free fictive prep was used as previously described^29^. Briefly, a small blade was used to scrape agarose away from the body up to the swim bladder, leaving the trunk and head of the fish immobilized while the tail could freely move.

#### 3.2. Zebrafish Calcium Imaging

OMR stimulus videos were generated using PsychoPy^52^ and cast onto a small piece of weigh paper (Fisherbrand, USA) placed below the dish using a projector (Sony, Japan). When the tail-free fictive prep was used, gross tail movements were recorded using a high-speed camera (Teledyne, USA) with a long-pass filter (λp ≥ 830 nm, Schott, Germany) positioned beneath the fish. Illumination was provided by an infrared LED array (Waveform Lighting, USA) adjacent to the objective above the fish. A small slit was cut in the weigh paper so fish tail movements were visible to the camera below. Both 10× and 20× objectives were immersed directly into the E3 media surrounding the fish. In both scenarios, imaging was carried out at 5 Hz volumetric rate, and a small region of interest was cropped on the camera chip to facilitate the necessary imaging speeds. After imaging, all fish were released into fresh E3 for recovery.

#### 3.3. Raw Data Processing

Raw image files from the microscope were preprocessed using custom MATLAB code to extract individual volumes and remove frames corresponding to the resetting of the galvo mirror. Additional correction for system shearing was calculated using an affine transform (affine3D, imwarp) with nearest neighbor interpolation.

#### 3.4. Anatomical Region Trace Extraction

For fish expressing cytosolic GCaMP imaged with the 10× objective, a single timepoint was used to map known anatomical regions to the imaging volumes. Registration to the MapZeBrain (MZB)^53^ reference atlas was performed using Advanced Normalization Tools (ANTs) as previously described^54,55^. Masks defining anatomical brain regions in the reference brain were downloaded from the MapZeBrain website (https://mapzebrain.org/) and manually curated for unique, non-overlapping regions that cumulatively covered the entire brain. Transformation matrices generated by ANTs were then inversely applied to each mask to register them to experimental volumes. Intensity values from across the entire ROI defined by the mask were averaged for each time point and plotted against time using custom MATLAB code. Cross-correlation analysis was done on these extracted traces using a custom R code (cor, corrplot). All images and mask overlays were produced manually using Fiji/ImageJ.

#### 3.5. Wavelet-Based Denoising with Dual-Stage Watershed Neuron Segmentation

For fish expressing nuclear localized GCaMP imaged with the 20× objective, a single timepoint was used for neuron segmentation. We applied Wavelet-Based Background and Noise Subtraction (WBNS) to remove low-frequency background and high-frequency noise^56^. The algorithm relies on multi-level discrete wavelet decomposition using the Haar wavelet, which separates each image into low-frequency approximation coefficients and high-frequency detail coefficients via sequential low-pass and high-pass filtering. Background was estimated by zeroing detail coefficients, followed by reconstruction with the inverse wavelet transformation and additional Gaussian smoothing. High-frequency noise was estimated from the finest detail level by setting the approximation coefficients to unity, retaining only the detail coefficients at the chosen noise level, and then reconstructing. Both background and noise images were subtracted sequentially from the original with a positivity constraint to yield the denoised signal. We implemented this process using the open-source WBNS Python package (https://github.com/NienhausLabKIT/HuepfelM/tree/master/WBNS/), with user-defined parameters for resolution and noise (resolution_px = 2, noise_lvl = 1).

After WBNS, binary masks were generated by global thresholding followed by removal of small objects (<15 voxels) to eliminate noise remnants and non-cellular debris. A two-stage watershed approach was then applied to improve object separation in densely packed regions. In the first stage, the 3D Euclidean distance transform was computed from the binary mask, and seeds were identified using a fast local-maxima detector (*scikit-image* implementation^57^) with a cubic footprint of 7×7×7 voxels. These seeds initiated the 3D marker-controlled watershed segmentation^57–59^. In the second stage, large components (>300 voxels) from the first segmentation, likely containing merged cells, were re-segmented using the same seeding method. Final labels were obtained by combining small components from the first stage with refined large components from the second, followed by morphological dilation (ball radius = 1 voxel) to smooth boundaries and ensure segmentation sizes matched neuronal dimensions. Alignment of the mask was verified in FIJI/ImageJ by manually defining ROIs for individual neurons and extracting the time course data (**Supplemental Fig. S5**)

#### 3.6. Cell Clustering

Clustering was completed using non-negative matrix factorization with 10 components on all individual neuron traces as previously described^35^. Neurons were assigned to the cluster in which they had the most significant contribution and then reorganized into clusters before plotting. Cluster assignments were then mapped back to the original mask by assigning a separate color to each individual cluster, and intensities were weighted by the relative contribution of the neuron to its respective cluster. Additionally, a weighted mask for each individual cluster was generated to identify anatomical regions where highly contributing neurons were located for that cluster. Fiji/ImageJ was used to generate representative images. Imaris was used to generate the 3D rendering of the mask.

### 4. Cleared Mouse Brain Imaging

#### 4.1. Brain Samples and Clearing Protocol

Four adult mouse brains were used in this study. For vasculature visualization, we used *Tie2-Cre;Ai9* double-transgenic animals (endothelial *Tek/Tie2* promoter–driven *Cre* crossed to *tdTomato* reporter), which yield *tdTomato* labeling of endothelial cells. A separate specimen was sparsely labeled for neurons by a retro-orbital injection of a dual-AAV cocktail driving tdTomato under an *hSyn-Cre/Tet-Off* design. Two additional specimens expressing Mobp-EGFP were stained for the Thy1 *CreERT2* cell line. Animals were maintained on a C57BL/6 background under approved protocols.

Two different clearing approaches were used in this study. In Route 1 (CUBIC), brains were perfusion-fixed in 4% PFA, washed in PBS at 37 °C, delipidated in CUBIC-L with periodic solution refreshes, and refractive-index-matched in CUBIC-R+ until samples were optically homogeneous. Samples were then secured in 30 mm dishes with UV-curable adhesive and immersed in the same matching medium^47^. In Route 2 (SHIELD → CUBIC-L → uRIMS), tissue was stabilized with SHIELD OFF/ON chemistry, delipidated in CUBIC-L, and finally index matched in a modified high RI uRIMS formulation tuned to either RI ≈ 1.496 or EasyIndex RI ≈ 1.52 at pH ≈ 7.4 to limit expansion^47,60,61^. When using high-RI RIMS/uRIMS, the bath was sealed with sunflower oil to prevent evaporation induced crystallization of the iodixanol solution. Both matching media were recovered after imaging, and any residual oil was removed before reusing the sample for additional imaging sessions. Additional reagent compositions, incubation schedules, and QC criteria are provided in the Supplemental Methods (**Supplemental Note 1**).

#### 4.2. Stage Triggered Acquisition

To avoid scan-induced aberrations, we disabled galvanometric scanning and formed volumetric images by moving the specimen using a pair of Thorlabs stages (MLJ250 for z-axis motion and Z825B for y-axis scanning). The y-axis stage controller (KDC101) generated one TTL pulse for each stage position, triggering the high-speed camera (Kinetix22, “Edge Trigger” mode) to collect one exposure per pulse. The complete scanning parameters for each magnification are provided in **Supplemental Tables 3 and 4**.

#### 4.3. Stitching and Post-processing

Postprocessing and stitching were performed in ImageJ (Fiji) and MATLAB R2024b using custom stitching code. The slices with the highest focal acuity were selected based on the orthogonal (x-z) or lateral (x-y) maximum projection views. Raw image stacks were cropped to the slice range, keeping ∼10% overlap between adjacent volumes. Each stack was converted to object coordinates with a slice-wise affine deskew transformation, implemented as a shear x ← x − s·z, with s derived from the oblique geometry at the corresponding magnification. Subsequent volumes were registered through a ROI-constrained procedure. A user defined rectangle on the terminal slice of the current volume was used as a template to identify the matching slice in the next volume as the maximum of the FFT-based cross-correlation from the mean subtracted frames. To compensate for slow magnification drift along the fast scan axis, the anisotropic scale was refined through a one-dimensional grid search over 0.95–1.00 with 0.001 increments to find the maximizer of the ROI correlation and degree of planar translation. We applied subpixel warping with bilinear interpolation based off the translation degree and blended across the boundary frames to suppress seams. Intensities were normalized to 16-bit using the global stack maximum. This lightweight, ROI guided FFT registration yields a seamless stitched full volume.

### 5. Sample preparation

All animal-related procedures including larval zebrafish, mouse whole-brain clearing and mouse *in vivo* imaging were in accordance with the Institutional Animal Care and Use Committee at Johns Hopkins University and conformed to the guidelines on the Use of Animals from the National Institutes of Health (NIH).

## Contribution

JY conceptualized the study; JY and DM setup the microscope; JY, HW implemented panoramic scanning protocol; DM performed *in vivo* mouse experiment; DM, JY analyzed blood flow data; DM, GK, HW performed *in vivo* Zebrafish larval experiment; GK, SL analyzed the Zebrafish data; HW setup the visual stimulus assay for larval zebrafish calcium imaging; LH performed resolution characterization analysis and Zemax simulation; YTKX performed optical clearing preparation; XL and JY performed imaging on optically cleared brain; XL and JY performed data analysis on cleared brain imaging data. DM, GK, XL, LH, JY prepared figures, and writing. JY, LT, JM, DEB supervised the study and coordinated the collaboration. JY supervised the study overall.

## Acknowledgement

This manuscript was supported in part by the National Institutes of Health (NIH, EB034272; AG072305; NS041435) and subject to the NIH Public Access Policy. The NIH has been given a right to make this manuscript publicly available in PubMed. We also acknowledge Boons Pickens endowment fund, Wilmer unrestricted grant from Research to Prevent Blindness, grant 2024-338490 from the Chan Zuckerberg Initiative DAF, an advised fund of Silicon Valley Community Foundation, as well as the Dr. Miriam and Sheldon G. Adelson Medical Research Foundation to partially support the study. YKTX and DM received support from a Kavli Neuroscience Discovery Institute Fellowship. GK received support from the Johns Hopkins Cross-Disciplinary Graduate Program in Biomedical Sciences. JSM received support from the Helen Larson and Charles Glenn Grover endowment funds. We thank Esther Whang for assistance with R based cross correlation analysis of zebrafish data.

## Notes

### Competing Interest Statement

The authors have declared no competing interest.

### Summary of Updates

This version has improved resolution in the PDF to correct some illegible text in the figures.

## References

1. Keller, P. J., Schmidt, A. D., Wittbrodt, J. & Stelzer, E. H. K. Reconstruction of Zebrafish Early Embryonic Development by Scanned Light Sheet Microscopy. Science 322, 1065–1069 (2008).

2. Tian, L. et al. Imaging neural activity in worms, flies and mice with improved GCaMP calcium indicators. Nat Methods 6, 875–881 (2009).

3. Glaser, A. K. et al. Light-sheet microscopy for slide-free non-destructive pathology of large clinical specimens. Nat Biomed Eng 1, 0084 (2017).

4. Kim, J. Recent advances in oblique plane microscopy. Nanophotonics 12, 2317–2334.

5. Dunsby, C. Optically sectioned imaging by oblique plane microscopy. Opt. Express 16, 20306 (2008).

6. Bouchard, M. B. et al. Swept confocally-aligned planar excitation (SCAPE) microscopy for high-speed volumetric imaging of behaving organisms. Nature Photon 9, 113–119 (2015).

7. Voleti, V. et al. Real-time volumetric microscopy of in vivo dynamics and large-scale samples with SCAPE 2.0. Nat Methods 16, 1054–1062 (2019).

8. Yang, B. et al. DaXi—high-resolution, large imaging volume and multi-view single-objective light-sheet microscopy. Nat Methods 19, 461–469 (2022).

9. Botcherby, E. J., Juškaitis, R., Booth, M. J. & Wilson, T. An optical technique for remote focusing in microscopy. Optics Communications 281, 880–887 (2008).

10. Bishop, K. W., Glaser, A. K. & Liu, J. T. C. Performance tradeoffs for single- and dual-objective open-top light-sheet microscope designs: a simulation-based analysis. Biomed. Opt. Express 11, 4627 (2020).

11. Alfred Millett-Sikking & Andrew York. High NA single-objective light-sheet. 10.5281/ZENODO.3376243 (2019) doi:10.5281/ZENODO.3376243.

12. Shao, W. et al. Mesoscopic oblique plane microscopy with a diffractive light-sheet for large-scale 4D cellular resolution imaging. Optica 9, 1374 (2022).

13. Daetwyler, S. et al. Mesoscopic oblique plane microscopy via light-sheet mirroring. Optica 10, 1571 (2023).

14. Lamb, J. R., Mestre, M. C., Lancaster, M. & Manton, J. D. Direct-view oblique plane microscopy. *Optica*, OPTICA 12, 469–472 (2025).

15. Singh, R. et al. Oblique plane microscope for mesoscopic imaging of freely moving organisms with cellular resolution. *Opt. Express*, OE 31, 2292–2301 (2023).

16. Yang, B. et al. Epi-illumination SPIM for volumetric imaging with high spatial-temporal resolution. Nat Methods 16, 501–504 (2019).

17. Hoffmann, M. & Judkewitz, B. Diffractive oblique plane microscopy. Optica 6, 1166 (2019).

18. Hoffmann, M., Henninger, J., Veith, J., Richter, L. & Judkewitz, B. Blazed oblique plane microscopy reveals scale-invariant inference of brain-wide population activity. Nat Commun 14, 8019 (2023).

19. Sapoznik, E. et al. A versatile oblique plane microscope for large-scale and high-resolution imaging of subcellular dynamics. eLife 9, e57681 (2020).

20. Marchand, P. J., Lu, X., Zhang, C. & Lesage, F. Validation of red blood cell flux and velocity estimations based on optical coherence tomography intensity fluctuations. Sci Rep 10, 19584 (2020).

21. Unekawa, M. et al. RBC velocities in single capillaries of mouse and rat brains are the same, despite 10-fold difference in body size. Brain Research 1320, 69–73 (2010).

22. Ogoh, S. Interaction between the respiratory system and cerebral blood flow regulation. Journal of Applied Physiology 127, 1197–1205 (2019).

23. Söderström, P. et al. Respiratory influence on cerebral blood flow and blood volume – A 4D flow MRI study. J Cereb Blood Flow Metab 45, 1531–1542 (2025).

24. Ewald, A. J., Werb, Z. & Egeblad, M. Monitoring of Vital Signs for Long-Term Survival of Mice under Anesthesia: FIGURE 1. Cold Spring Harb Protoc 2011, pdb.prot5563 (2011).

25. Rock, I. & Smith, D. The Optomotor Response and Induced Motion of the Self. Perception 15, 497–502 (1986).

26. Neuhauss, S. C. F. et al. Genetic Disorders of Vision Revealed by a Behavioral Screen of 400 Essential Loci in Zebrafish. J. Neurosci. 19, 8603–8615 (1999).

27. Orger, M. B., Smear, M. C., Anstis, S. M. & Baier, H. Perception of Fourier and non-Fourier motion by larval zebrafish. Nat Neurosci 3, 1128–1133 (2000).

28. Portugues, R. & Engert, F. The neural basis of visual behaviors in the larval zebrafish. Current Opinion in Neurobiology 19, 644–647 (2009).

29. Jouary, A., Haudrechy, M., Candelier, R. & Sumbre, G. A 2D virtual reality system for visual goal-driven navigation in zebrafish larvae. Sci Rep 6, 34015 (2016).

30. Roeser, T. & Baier, H. Visuomotor Behaviors in Larval Zebrafish after GFP-Guided Laser Ablation of the Optic Tectum. J. Neurosci. 23, 3726–3734 (2003).

31. Nikolaou, N. et al. Parametric Functional Maps of Visual Inputs to the Tectum. Neuron 76, 317–324 (2012).

32. Gabriel, J. P., Trivedi, C. A., Maurer, C. M., Ryu, S. & Bollmann, J. H. Layer-Specific Targeting of Direction-Selective Neurons in the Zebrafish Optic Tectum. Neuron 76, 1147–1160 (2012).

33. Robles, E., Laurell, E. & Baier, H. The Retinal Projectome Reveals Brain-Area-Specific Visual Representations Generated by Ganglion Cell Diversity. Current Biology 24, 2085–2096 (2014).

34. Kramer, A., Wu, Y., Baier, H. & Kubo, F. Neuronal Architecture of a Visual Center that Processes Optic Flow. Neuron 103, 118–132.e7 (2019).

35. Mu, Y. et al. Glia Accumulate Evidence that Actions Are Futile and Suppress Unsuccessful Behavior. Cell 178, 27–43.e19 (2019).

36. Chen, X. et al. Brain-wide Organization of Neuronal Activity and Convergent Sensorimotor Transformations in Larval Zebrafish. Neuron 100, 876–890.e5 (2018).

37. Naumann, E. A. et al. From Whole-Brain Data to Functional Circuit Models: The Zebrafish Optomotor Response. Cell 167, 947–960.e20 (2016).

38. Costantini, I., Cicchi, R., Silvestri, L., Vanzi, F. & Pavone, F. S. In-vivo and ex-vivo optical clearing methods for biological tissues: review. Biomed. Opt. Express 10, 5251 (2019).

39. Ueda, H. R. et al. Tissue clearing and its applications in neuroscience. Nat Rev Neurosci 21, 61–79 (2020).

40. Chen, F., Tillberg, P. W. & Boyden, E. S. Expansion microscopy. Science 347, 543–548 (2015).

41. Principles of Neural Science. (McGraw-Hill medical, New York, 2013).

42. Mandelli, M. L. et al. Diffusion Tensor Imaging of Spinocerebellar Ataxias Types 1 and 2. American Journal of Neuroradiology 28, 1996–2000 (2007).

43. Fujishima, K., Kurisu, J., Yamada, M. & Kengaku, M. βIII spectrin controls the planarity of Purkinje cell dendrites by modulating perpendicular axon-dendrite interactions. Development 147, dev194530 (2020).

44. Prakash, N. et al. Patterns of fractional anisotropy changes in white matter of cerebellar peduncles distinguish spinocerebellar ataxia-1 from multiple system atrophy and other ataxia syndromes. NeuroImage 47, T72–T81 (2009).

45. Gao, Y. et al. β-III Spectrin Is Critical for Development of Purkinje Cell Dendritic Tree and Spine Morphogenesis. J. Neurosci. 31, 16581–16590 (2011).

46. Nicoletti, G. et al. MR Imaging of Middle Cerebellar Peduncle Width: Differentiation of Multiple System Atrophy from Parkinson Disease. Radiology 239, 825–830 (2006).

47. Susaki, E. A. et al. Versatile whole-organ/body staining and imaging based on electrolyte-gel properties of biological tissues. Nat Commun 11, 1982 (2020).

48. Shao, W. et al. Wide field-of-view volumetric imaging by a mesoscopic scanning oblique plane microscopy with switchable objective lenses. Quant Imaging Med Surg 11, 983–997 (2020).

49. Alfred Millett-Sikking, Any immersion remote refocus (AIRR) microscopy. Preprint at 10.5281/ZENODO.7425649 (2022).

50. Zhao, Y. et al. Isotropic super-resolution light-sheet microscopy of dynamic intracellular structures at subsecond timescales. Nat Methods 19, 359–369 (2022).

51. White, R. M., et al. Transparent Adult Zebrafish as a Tool for In Vivo Transplantation Analysis, Cell Stem Cell 2, 183–189 (2008).

52. Peirce, J. et al. PsychoPy2: Experiments in behavior made easy. Behav Res 51, 195–203 (2019).

53. Shainer, I. et al. A single-cell resolution gene expression atlas of the larval zebrafish brain. Sci. Adv. 9, eade9909 (2023).

54. Marquart, G. D. et al. High-precision registration between zebrafish brain atlases using symmetric diffeomorphic normalization. GigaScience 6, gix056 (2017).

55. Kunst, M. et al. A Cellular-Resolution Atlas of the Larval Zebrafish Brain. Neuron 103, 21–38.e5 (2019).

56. Hüpfel, M., Yu. Kobitski, A., Zhang, W. & Nienhaus, G. U. Wavelet-based background and noise subtraction for fluorescence microscopy images. Biomed. Opt. Express 12, 969 (2021).

57. Van Der Walt, S. et al. scikit-image: image processing in Python. PeerJ 2, e453 (2014).

58. Vincent, L. & Soille, P. Watersheds in digital spaces: an efficient algorithm based on immersion simulations. IEEE Trans. Pattern Anal. Machine Intell. 13, 583–598 (1991).

59. Soille, P. J. & Ansoult, M. M. Automated basin delineation from digital elevation models using mathematical morphology. Signal Processing 20, 171–182 (1990).

60. Swaney, J. et al. Scalable image processing techniques for quantitative analysis of volumetric biological images from light-sheet microscopy. Preprint at 10.1101/576595 (2019).

61. Takahashi, K., Kubota, S. I., Ehata, S., Ueda, H. R. & Miyazono, K. Protocol for Imaging and Analysis of Mouse Tumor Models with CUBIC Tissue Clearing. STAR Protocols 1, 100191 (2020).

