## Supplemental Materials for "Versatile high-speed volumetric imaging from microscopic to macroscopic scale by self-adaptive oblique plane microscopy"

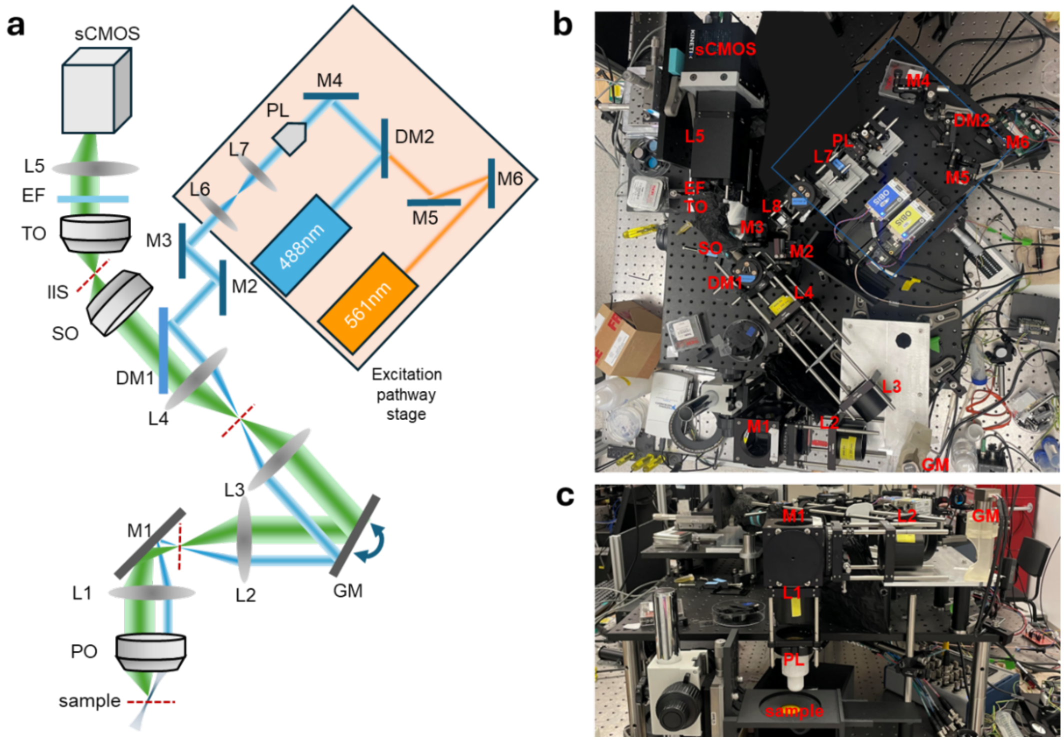


**Supplemental Figure S1. System construction. (a)** Layout of overall system with labeled components. The excitation paths include the 561 nm pathway (orange) which is steered by two mirrors (M6 and M5). This pathway combines pathways with the 488 nm pathway (blue) after passing through DM2, which reflects the 488 nm pathway. M4 redirects the excitation pathway to the Powell Lens (PL: 60° fan angle) which generates a static light sheet. L7 (f=30 mm) and L6 (f=100 mm) form a telescope to increase the light sheet width. M3 and M2 provide additional beam steering capabilities. These upstream components are all mounted on the same linear translation stage. DM1 reflects the excitation path to the L4 and L3 (both f=100 mm, 2xAC508-200-A) relay system. The galvanometer (GM) scans the beam across the y range during imaging. L2 and L1 (both f=100 mm, 2xAC508-200-A) form a final relay system. M1 changes the propagation direction from horizontal to vertical so that it travels downwards through the primary objective (PO) for upright sample mounting geometry. The position of the excitation pathway is fine-tuned with the excitation pathway linear translation stage to maximize the degree of lateral offset in the back pupil of the primary objective (PO). The PO focuses the excitation light-sheet at an angle onto the sample, completing the illumination path. The PO then collects the fluorescence signal emitted from this obliquely illuminated (titled) plane, utilizing the full aperture to capture the resulting wide cone of light. The emission pathway shares many of the initial optics with the excitation pathway (L1+L2 relay, M1, GM, and L3+L4 relay). Due to this shared, synchronized geometry, the emission is simultaneously de-scanned to produce a stationary downstream image. The secondary objective (SO) re-images the sample at the intermediate imaging space (IIS) after passing through DM1. This intermediate image retains the tilted focal plane carried over from the oblique sample plane. To create a planar final image that matches the detector, a remote imaging system (RIS) comprised of a tertiary objective (TO), emission filter (EF), tube lens (L5, f=120 mm), and sCMOS detector (Kinetix22) is rotated along the optical axis of the SO. The axial position of the RIS can be finely adjusted using a linear stage to ensure the detector plane perfectly matches the now-planar final image projected by the RIS. The actual system is shown with a top-down view **(b)** and side view **(c)**.

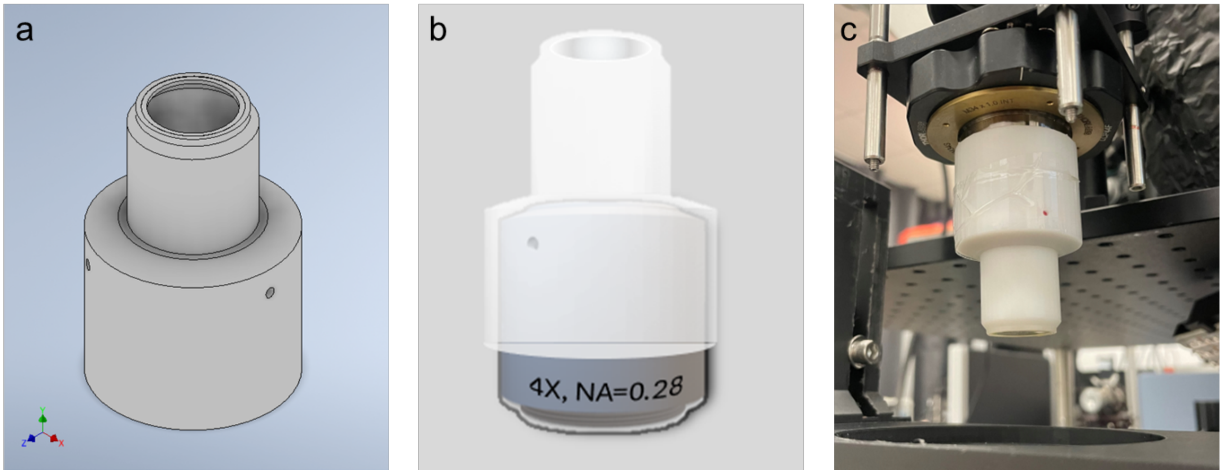


**Supplemental Figure S2. Liquid immersion cap for 4× objective. (a)** 3D design of liquid immersion cap for 4× objective during imaging. A glass cover slip was glued with epoxy to the cap opening. **(b)** The cap slides onto the tip of the objective so that it is flush with the objective lens barrel. This cap design maintains an air chamber with a constant depth around the tip of the objective, with the outer edge of the cover slip positioned at the working distance of the objective (WD=29 mm). **(c)** The objective with the cap was mounted in place with a magnetic adapter.

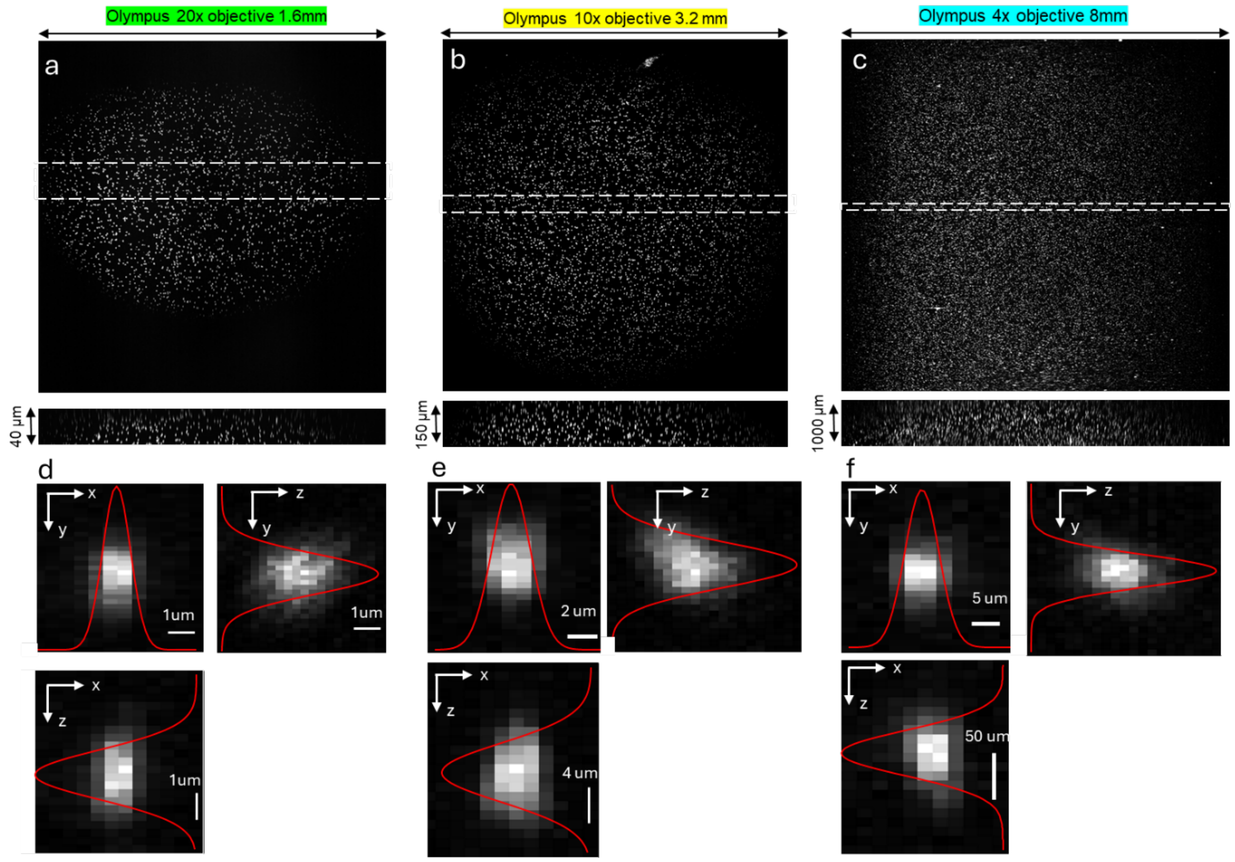


**Supplementary Figure S3. Point spread function across the FOV.** Images of fluorescent microspheres in 0.1% agarose were captured across the full FOV range with optical scanning by the galvanometer for (a) 20×, (b) 10×, (c) 4× Olympus objective. The x-z cross-sectional images show the maximum intensity projection over the dashed regions to demonstrate the depth of view. (d-f) Example of single isolated beads from the center of the FOV. Maximum intensity projections are shown in three planes. The Gaussian fitting was performed from the maximum projection along each axis, and the full-width-half-maximum was used to characterize the resolution for each dimension.

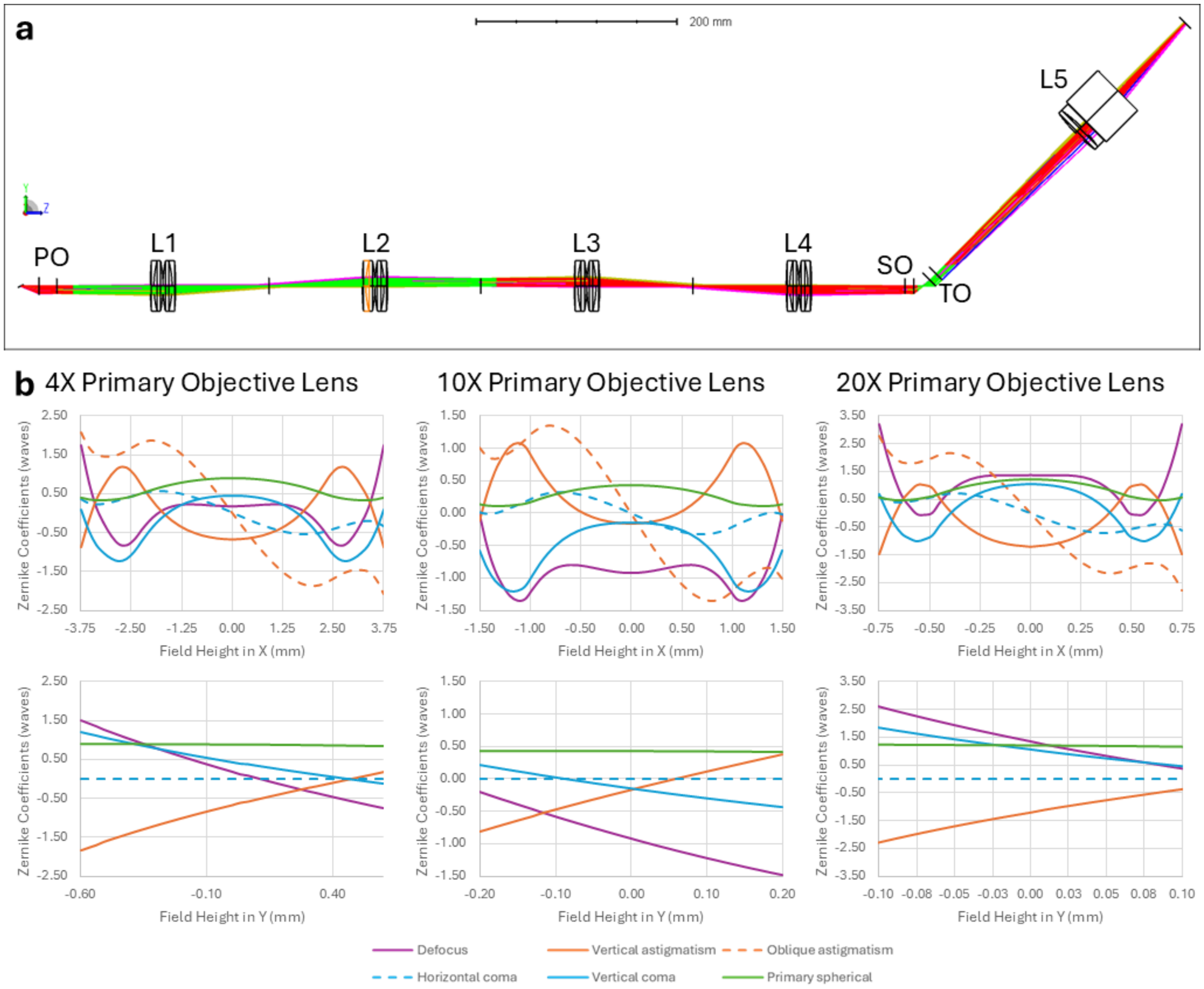


**Supplemental Figure S4. Theoretical optical aberration simulation.** (a) Layout of Zemax simulation of our system. We used lens models from the manufacturers and applied the paraxial lens approximation for the three objectives to evaluate aberrations caused by the relay optics and tertiary imaging system. Lens placements were optimized for smallest spot RMS with the 10× objective. **(b)** Aberration variance over the field of view for three different primary objectives (Olympus 4x, NA=0.28; Olympus 10×, NA=0.6; Olympus 20×, NA=1.0). The Zernike aberrations follow an M-shaped pattern across the FOV in the X direction similar to the fluorescent bead FWHM pattern in **Fig. 1i**.

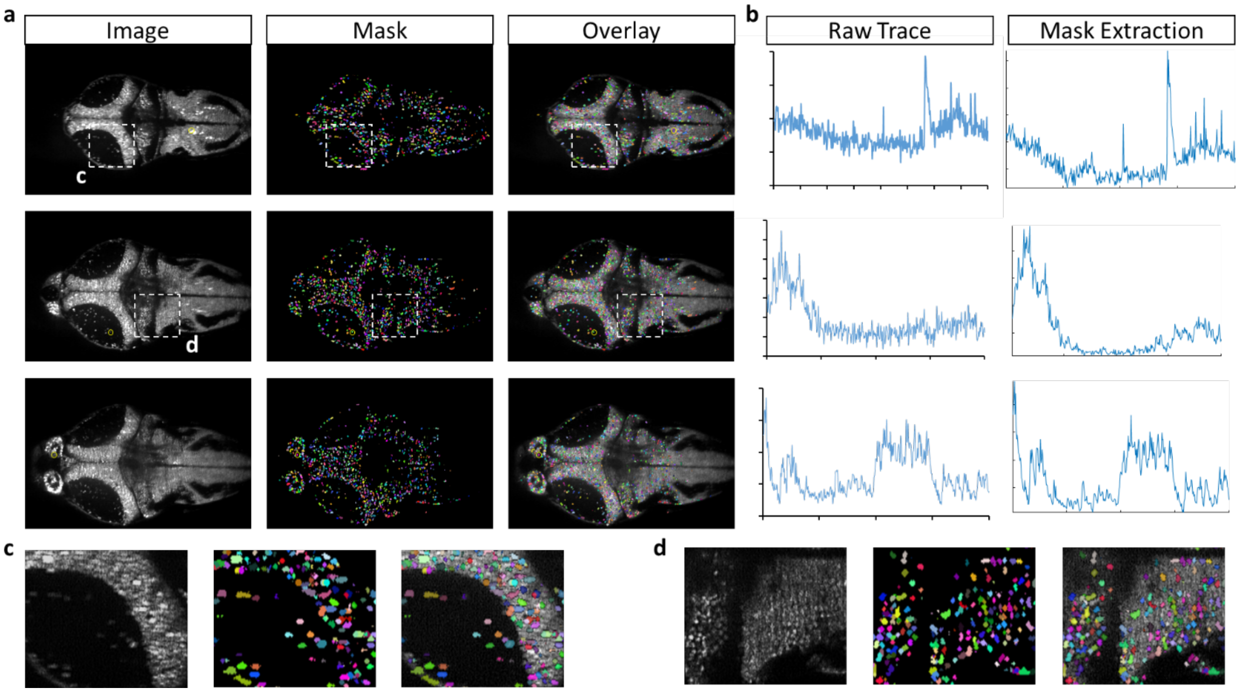


**Supplemental Figure S5. Verification of Single Neuron Segmentation. (a)** Single slices from imaging data (same as **Fig. 3g-i**), left; the color-coded segmentation mask, middle; and mask overlaid on image, right. **(b)** Intensity traces over the entire imaging time course for individual neurons (location indicated by the yellow circles in **(a)** extracted using a manually drawn ROI in FIJI, left; or by averaging all pixels corresponding to the specific neuron as defined by the mask, right. **(c, d)** Insets from **(a)** showing a zoomed in view of the imaging data, segmentation mask, and overlay.

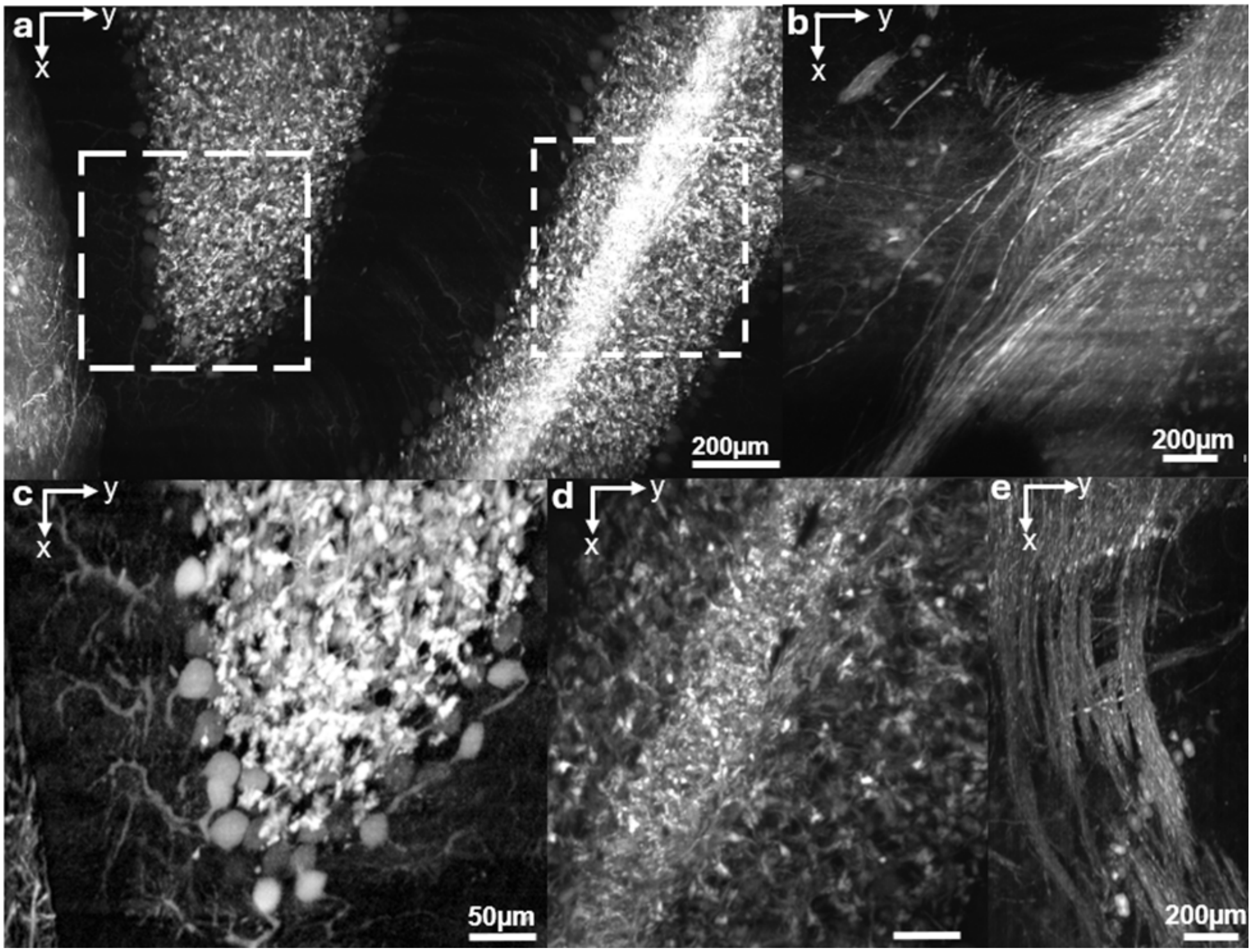


**Supplementary Figure S6. Scanning Over Cerebellum.** **(a)** 20× acquisition from the cerebellum of a Thy1 Slick H–YFP mouse (RI = 1.52). Maximum intensity projection over 0.08 mm depth shows the trilaminar organization; dashed boxes denote ROIs. **(b)** 20× maximum intensity projection over 0.08 mm at a shallower slice of the cerebellar peduncles, revealing bundled axons. **(c)** Zoom of ROI 1 from **(a)**, contrast adjusted to highlight single Purkinje cell bodies and adjacent granule cells. **(d)** Zoom of ROI 2 from **(a)** showing granule and molecular layers with mossy fibers. **(e)** 20× MIP at the deepest peduncle level demonstrating tightly packed, aligned fibers. Scale bars: 200 µm in **(a,b,e)**, 50 µm in **(c,d)**. Complete acquisition scheme is detailed in **Supplementary Note 2.**

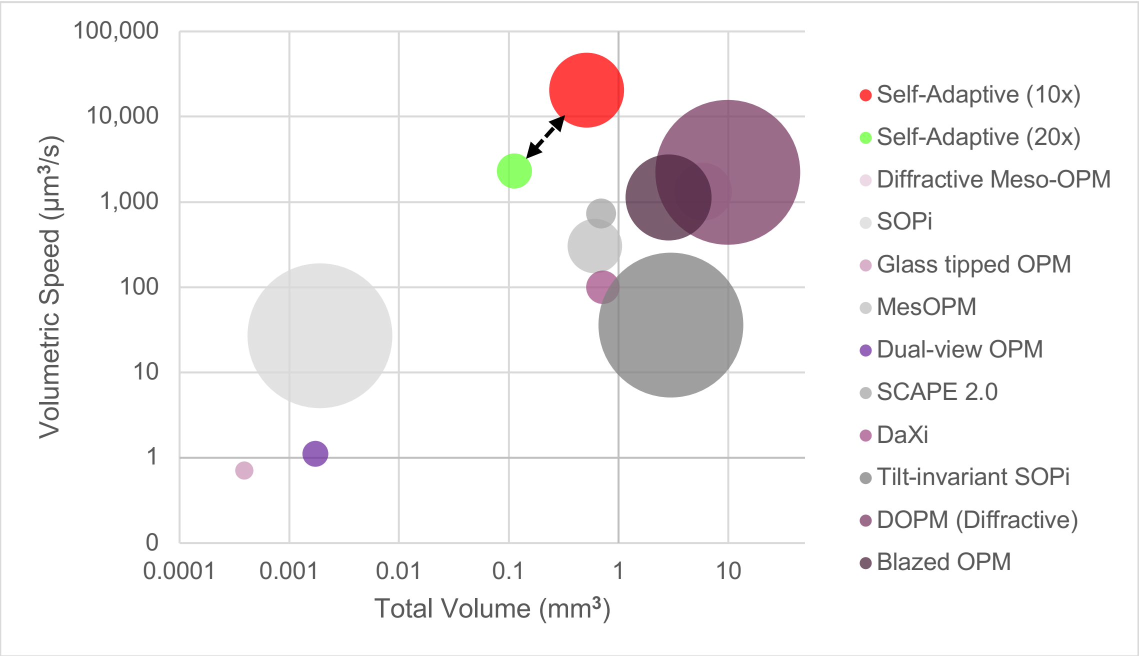


**Supplemental Figure S7. Volumetric Performance Comparison of Single Objective Light Sheet Microscopy Systems.** Volumetric speed ($\mu m^3/s$) is plotted against the total volume acquired ($mm^3$) on log scales. The marker size is scaled proportionally to the axial resolution where smaller markers indicate better resolution. The 10× Mesoscopic Self-Adaptive OPM (red) achieves the highest volumetric throughput across all systems without sacrificing the axial resolution when compared to other traditional mesoscopic single objective systems (gray markers). Its unique performance envelope (dashed black line) connects the mesoscopic mode with its 20× high-resolution microscopic mode (green). This higher resolution mode maintains axial resolution and volumetric throughput compared to other systems with similar axial resolution, at the cost of a slightly narrower FOV, without added significant optical complexity (pink and purple markers).

**Supplemental Table 1: Performance Comparison across State-of-the-art Single Objective Light Sheet Microscopy Systems.**

Performance comparison across premiere high-speed, large FOV, single objective light sheet systems shows that our system (Self-Adaptive 10×/20×) achieves superior volumetric throughput over other competitors. The total volume for each system was taken from reported values. The volumetric speed for each system was calculated using a dynamic imaging case from their respective published capability by multiplying the volume rate over the specified imaging volume.

| System Name | X Res (μm) | Y Res (μm) | Z Res (μm) | Voxel Volume ($\mu m^3$) | Volume Rate (Hz) | Total Volume ($mm^3$) | Volumetric Speed ($mm^3/s$) |
|---|---|---|---|---|---|---|---|
| **Self-Adaptive (10×)** | ***3*** | ***3*** | ***10*** | ***90*** | ***10*** | ***0.512*** | ***20.48*** |
| **Self-Adaptive (20×)** | ***1.2*** | ***2*** | ***2.2*** | ***5.28*** | ***100*** | ***0.113*** | ***2.31*** |
| Diffractive Meso-OPM[1] | *2.5* | *3* | *6* | *45* | *5* | *5.88* | *1.32* |
| SOPi[2] | *1.3* | *1.3* | *37.4* | *63.21* | *10* | *0.0019* | *0.027* |
| Glass tipped OPM[3] | *0.3* | *0.34* | *0.59* | *0.06* | *10* | *0.00039* | *0.00071* |
| MesOPM[3] | *1.5* | *2.8* | *5.3* | *22.26* | *0.5* | *0.61* | *0.304* |
| Dual-view OPM[4] | *0.46* | *0.59* | *1.2* | *0.33* | *2* | *0.0017* | *0.0011* |
| SCAPE 2.0[5] | *0.6* | *1.2* | *1.6* | *1.15* | *300* | *0.69* | *0.73* |
| DaXi[6] | *0.45* | *0.45* | *2* | *0.41* | *2* | *0.72* | *0.1* |
| Tilt-invariant SOPi[7] | *2.6* | *3.1* | *37.4* | *0.03* | *10* | *3.00* | *0.036* |
| Original OPM[8] | *0.82* | *0.82* | *4.3* | *2.89* | *1* | *0.00076* | *X* |
| DOPM (Diffractive)[9] | *2.6* | *3.1* | *37.4* | *301.44* | *25* | *9.90* | *2.23* |
| Blazed OPM[10] | *2.8* | *2.4* | *13.2* | *88.7* | *10* | *2.86* | *1.13* |

**Supplementary Table 2: Part labels for all optical components in Meso-OPM.**

Part names correspond with labels in **Supplemental Figure S1.**

| Component | Part Number | Focal length | Other Specs |
|---|---|---|---|
| PO – 20× | *XLUMPlanFL N* | *9 mm* | *NA=1.0, WD=12mm* |
| PO – 20× | *Clr Plan-Neofluar* | *8.25mm* | *NA=1.0, WD=6.4mm* |
| PO – 10× | *XLUMPlanFl* | *18 mm* | *NA=0.6, WD=3mm* |
| PO – 4× | *XLFLUOR4X-2/340* | *45 mm* | *NA=0.28, WD=29.5mm* |
| L1 | *2xAC508-200-A* | *100 mm* | |
| M1 | *PF10-03-P01* | | |
| L2 | *2xAC508-200-A* | *100 mm* | |
| Galvanometer | *Nutfield QS-12 OPD* | | *20mm aperture* |
| L3 | *2xAC508-200-A* | *100 mm* | |
| L4 | *2xAC508-200-A* | *100 mm* | |
| DM1 | *Di01-R405/488/561/635/800-t3-25x36* | | *IDEX Semrock* |
| SO | *20X: XLUMPlanFL N* | | |
| TO | *20X: XLUMPlanFL N* | | |
| EF | *ET535/70* | | |
| L5 | *AC508-300-AB + TTL200MP* | *120 mm* | |
| sCMOS Camera | *Kinetix* | | *15.6x15.6mm FOV, 5.7 MP, 664 fps, 6.5µm pixel size, 95% QE, -0.7$e^{-}$ read noise* |
| M2 | *PFR10-P01* | | |
| M3 | *UM10-AG* | | |
| L6 | *AC254-100-AB* | *100 mm* | |
| L7 | *AC254-030-AB* | *30 mm* | |
| PL | *LGL160* | | *60° fan angle* |
| M4 | *PF10-03-P01* | | |
| DM2 | *DMLP505R* | | *Long pass, 506nm cut on wavelength (or 495nm)?* |
| LS1 | *Coherent OBIS 488nm* | | |
| M5 | *PF10-03-P01* | | |
| M6 | *PF10-03-P01* | | |
| LS2 | *Coherent OBIS 561nm* | | |
| DC Power supply | *Siglent SPD3303X* | | *2x 32V/3.2A, 2.5-5V/3.2A, 1mV* |
| DAQmx board | *NI USB-6363* | | |
| Manual stage | | | |
| Motorized stage | *PT1-Z825B* | | *25 mm of travel distance* |
| Stage driver | *KDC101* | | |

**Supplemental Table 3. Imaging protocols sorted by biological application.**

Larval zebrafish calcium imaging

| Objective | Number of Rows | Frame Rate | Exposure Time | Camera Mode | Laser Power | Power at Sample |
|---|---|---|---|---|---|---|
| 10× | *250* | *~1133 fps* | *883 µs* | *Edge Trigger* | *488nm, 8 mW* | *0.97 mW* |
| 20× | *300* | *~944 fps* | *1059 µs* | *Edge Trigger* | *488nm, 8 mW* | *0.32 mW* |

Mouse *in vivo* cerebrovascular imaging

| Objective | Number of Rows | Frame Rate | Exposure Time | Camera Mode | Laser Power | Power at Sample |
|---|---|---|---|---|---|---|
| 10× | *150* | *~2667 fps* | *100 µs* | *Trigger First* | *488nm, 20 mW* | *4.84 mW* |
| 10× (rolling window) | *100* | *~9524 fps* | *100 µs* | *Edge Trigger* | *488nm, 20 mW* | *4.84 mW* |
| 20× | *150* | *~7271 fps* | *100 µs* | *Trigger First* | *488nm, 20 mW* | *1.27 mW* |

Cleared whole mouse brain imaging

| Objective | Number of Rows | Frame Rate | Exposure Time | Camera Mode | Laser Power | Power at Sample |
|---|---|---|---|---|---|---|
| 4× | *300* | *~89.9 fps* | *1125 µs* | *Edge Trigger* | *488 nm, 10mW* | *3.19 mW* |
| | | | | | *488nm, 20 mW* | *6.23 mW* |
| | | | | | *561 nm, 45 mW* | *16.45 mW* |
| 10× | *300* | *~89.9 fps* | *1125 µs* | *Edge Trigger* | *488 nm, 10 mW* | *2.31 mW* |
| | | | | | *561 nm, 10 mW* | *2.81 mW* |
| 20× | *300* | *~89.9 fps* | *1125 µs* | *Edge Trigger* | *488 nm, 5 mW* | *0.51 mW* |
| | | | | | *561 m, 5 mW* | *0.46 mW* |

**Supplemental Table 4: Stage-Triggered Scanning Parameters.**

Image volumes were acquired by translating the specimen along the y-axis and triggering the camera exposure for one frame per stage generated TTL pulse. The Thorlab stage Kinesis GUI was set with I/O Mode to 'OUT - At Position Steps (Fwd)' and I/O Polarity to 'Trigger High', so that the controller outputs a TTL pulse at each forward step to trigger the camera. The Teledyne Kinetix22 camera was run in the dynamic range (16 bit) acquisition mode with external edge triggering and an ROI width of 3200 columns. Readout speed was 3.749 μs/line. The trigger interval, trigger count, and start position were specified per acquisition to match the desired scan extent. To limit motion blur and satisfy the light sheet confocal sampling condition, the camera sensor was windowed to the central 300 rows. The exposure time was set based on sample intensity (typically 5–15 ms) so that the signal remained within the dynamic range of the detector for different laser powers, magnifications, and samples. The duration of a single frame was calculated by adding the camera readout overhead to the exposure duration. The trigger frequency was set so that one image was acquired for each frame interval (i.e., one trigger per exposure-plus-readout period). The stage speed was then chosen so that the stage advanced by exactly one axial step during each frame interval. For each magnification, the step size was selected to satisfy Nyquist sampling and avoid under sampling. Magnification specific settings (exposure, step size, and stage speed) are summarized below.

| Objective/ ROI | Y Sanning Range | Number of Frames | Exposure & Readout Time | Stage Velocity | Z Depth Step Size |
|---|---|---|---|---|---|
| 4× Whole Brain | *12.5 -15.5 mm* | *4000* | *11.5 ms* | *0.349 -0.562 mm/s* | *0.75 mm* |
| 4× Regional | *2 - 4 mm* | *2000 - 4000* | *6.5 ms -16.5 ms* | *0.061 - 0.153 mm/s* | *0.75 mm* |
| 10× Mesoscale | *2 - 4mm* | *2000 - 4000* | *11.5 ms* | *0.045 - 0.09 mm/s* | *0.2 mm* |
| 20× Single Cell | *2mm* | *4000* | *11.5 ms* | *0.09 mm/s* | *0.05 mm* |

**Supplementary Note 1. Cleared Brain Protocol.**

**Animal care and use**

All animal experiments were conducted in accordance with protocols approved by the Animal Care and Use Committee at Johns Hopkins University. Both female and male adult mice were used and randomly assigned to experimental groups. All animals were healthy and exhibited no overt behavioral abnormalities. Generation and genotyping of BAC transgenic lines from *Mobp-GFP* (GENSAT[11]) have been described previously[12]. Mice were housed in a climate-controlled room on a 12-hour light/dark cycle, in groups of no more than five per cage. Food and water were provided ad libitum, except during cuprizone administration. All animals were maintained on a pure C57BL/6 (B6) background. Other mice included *Mobp-EGFP* crossed with *Tie2-Cre;Ai9* (Jackson laboratory) reporter animals to visualize the brain vasculature.

**AAV injections for sparse viral labeling**

To achieve sparse and inducible labeling of neuronal populations, 21-day-old mice were injected retro-orbitally with a mixture of adeno-associated viruses (AAVs). The mixture contained three viruses: *AAV-PHP.eB-Syn-Flex-TREx2-tTA* ($3.40 \times 10^{13}$ genome copies [GC]/mL), *AAV-PHP.eB-7xTRE-tdTomato* ($4.84 \times 10^{13}$ GC/mL), and *AAV9-hSyn-Cre*. Viruses were combined at a 1:100 volume ratio and diluted in sterile saline to a final injection volume of 50 µL per mouse. Injections were performed under brief isoflurane anesthesia (1.5%) following induction at 5%. Virus was delivered slowly into the retro-orbital sinus over 5–10 seconds to minimize reflux, and gentle pressure was applied to the eyelid after injection. Mice were monitored during recovery and returned to their home cage once normal activity resumed. Animals were perfused three weeks later with 4% paraformaldehyde (PFA). All viruses were acquired from Janelia research campus. We achieved sparse viral labeling of random neurons by injecting a cocktail of AAVs through the retro-orbital sinus (*AAV-hSyn-Cre; AAV-Syn-Flex-TREx2-tTA; AAV-7xTRE-tdTomato*). The *hSyn* driver ensured neuron-specificity while the tet-off system amplified tdTomato expression once a neuron had been infected with all three viruses.

**Cleaning Route 1: CUBIC L to CUBIC R+**

Brains were perfusion fixed with 4% paraformaldehyde, post fixed overnight at 4° C, and washed three times in PBS at 37°C with gentle shaking to remove residual fixative. Delipidation proceeded in CUBIC L with a brief equilibration in 50% CUBIC L at 37°C (≥6h) followed by incubation in 100% CUBIC L at 37°C for 1-7 days. Solution was refreshed every 3-4 days and vessels were kept covered to minimize evaporation; endpoint was defined by loss of color in the bath and visual translucency of the brain. For consistency across animals, we recorded wet mass and linear dimensions before delipidation and after each solution change to quantify expansion or shrinkage. When immunolabeling was required, delipidation was followed by antibody incubation in detergent containing buffer at 34°C for 7 days with agitation and light protection, then thorough PBS washes.

Refractive index matching was performed in CUBIC R+ by stepping samples from 50% solution at room temperature in the dark (6 h) to 100% solution (≥12h until optical homogeneity). All solutions were degassed prior to use to reduce microbubble formation in deep tissue. For mounting, cleared brains were transferred to a 30 mm dish and secured to the base with a small fillet of UV curable adhesive applied away from the imaging field; the dish was then filled with RIMS to cover the sample completely. If buoyancy or residual tissue softness caused motion during stage sweeps, we stabilized the preparation by embedding the sample in a thin bed of low melting agarose prepared in CUBIC R and allowed to gel at room temperature before imaging. Adhesives, plastics, and seals in contact with the bath were chosen for compatibility with hydrophilic media; we avoided materials that leach plasticizers or fluoresce under 488/561 nm excitation. Samples were stored at 4°C in fresh RIMS prior to imaging, protected from light, and re checked for RI drift immediately before acquisition.

**Clearing Route 2: SHIELD → CUBIC-L → uRIMS**

Brains were perfusion fixed with 4% paraformaldehyde, post fixed overnight at 4°C, and washed three times in PBS at 37°C with gentle shaking to remove residual fixative. Transferred from PBS directly into SHIELD OFF at 4 °C (50% SHIELD buffer, 25% SHIELD Epoxy in $dH_2O$) with gentle nutation and light protection for 3 days, then moved to SHIELD ON for overnight incubation at 37 °C to complete epoxide crosslinking. Following SHIELD, tissue was rinsed in PBS for 1 h at room temperature and then overnight at 37°C. Delipidation was performed in 100% CUBIC-L at 37°C with solution replacement every 3–4 days; the duration scaled with tissue age and lipid content, and we noted that SHIELD-processed samples can appear more opaque than CUBIC-only brains despite remaining suitable for deep imaging. Brains were washed in PBS for 1 h at room temperature and then overnight at 37°C to remove residual detergents.

Final matching used a urea-augmented high-RI medium (uRIMS) adjusted to RI ≈ 1.496 at pH ≈ 7.4. The working formulation comprised ~60% w/v Nycodenz, 0.5% w/v meglumine, 0.25% w/v diatrizoic acid, and 0.01% sodium azide in 20 mM phosphate buffer, supplemented with 40% w/v urea; Tween-20 was omitted to prevent expansion and bubble formation. Solutions were prepared fresh, filtered, degassed, and verified on a refractometer before use. Samples typically reached near-transparent appearance within ~1–2 days at 37 °C with gentle agitation. Increasing RI beyond the target by adding Nycodenz did not improve image quality and increased bath turbulence near saturation; we therefore maintained the target RI and controlled pH at ~7.4 to limit tissue expansion while preserving EGFP signal. Long-term storage in uRIMS was avoided because the aqueous, high-urea environment gradually softens tissue; instead, brains were briefly rinsed, stored dry in airtight tubes at room temperature, and re-equilibrated in fresh uRIMS overnight at 37°C before imaging.

**Supplementary Note 2. Imaging protocol for cerebellar imaging.**

Cerebellar imaging (**Fig. S6**) was performed in an adult Thy1-Slick-H–YFP mouse cleared with Route 2 imaged in EasyIndex at RI = 1.52. The same region was acquired at 10× and 20× to resolve the trilaminar architecture (molecular layer, Purkinje cell layer with planar dendritic arbors, and granule cell layer) and to delineate cerebellar peduncles.

A single field covering 1.5 mm×2mm at the cerebellar hemisphere was imaged with a depth stratified plan: 12 volumes with z = 0.2 mm in 10× magnification and 40 volumes in 20× magnification with z = 0.05 mm. The first 20 volumes sampled the superficial laminae (molecular→Purkinje→granule). We then skipped 0.4 mm to avoid repetitive information and acquired the remaining 20 volumes at the deepest accessible depth in this ROI to capture the cerebellar peduncles.

**References:**


1. Shao, W. *et al.* Mesoscopic oblique plane microscopy with a diffractive light-sheet for large-scale 4D cellular resolution imaging. *Optica* **9**, 1374 (2022).

2. Sapoznik, E. *et al.* A versatile oblique plane microscope for large-scale and high-resolution imaging of subcellular dynamics. *eLife* **9**, e57681 (2020).

3. Singh, R. *et al.* Oblique plane microscope for mesoscopic imaging of freely moving organisms with cellular resolution. *Opt. Express, OE* **31**, 2292–2301 (2023).

4. Sparks, H. *et al.* Dual-view oblique plane microscopy (dOPM). *Biomed. Opt. Express* **11**, 7204 (2020).

5. Voleti, V. *et al.* Real-time volumetric microscopy of in vivo dynamics and large-scale samples with SCAPE 2.0. *Nat Methods* **16**, 1054–1062 (2019).

6. Yang, B. *et al.* DaXi—high-resolution, large imaging volume and multi-view single-objective light-sheet microscopy. *Nat Methods* **19**, 461–469 (2022).

7. Kumar, M. & Kozorovitskiy, Y. Tilt-invariant scanned oblique plane illumination microscopy for large-scale volumetric imaging. *Opt. Lett.* **44**, 1706 (2019).

8. Dunsby, C. Optically sectioned imaging by oblique plane microscopy. *Opt. Express* **16**, 20306 (2008).

9. Hoffmann, M. & Judkewitz, B. Diffractive oblique plane microscopy. *Optica* **6**, 1166 (2019).

10. Hoffmann, M., Henninger, J., Veith, J., Richter, L. & Judkewitz, B. Blazed oblique plane microscopy reveals scale-invariant inference of brain-wide population activity. *Nat Commun* **14**, 8019 (2023).

11. Gong, S. *et al.* A gene expression atlas of the central nervous system based on bacterial artificial chromosomes. *Nature* **425**, 917–925 (2003).

12. Hughes, E. G., Orthmann-Murphy, J. L., Langseth, A. J. & Bergles, D. E. Myelin remodeling through experience-dependent oligodendrogenesis in the adult somatosensory cortex. *Nat Neurosci* **21**, 696–706 (2018).
